# Rapid repurposing of microvillar content drives a flagellate-to-amoeboid switch in the closest relative of animals

**DOI:** 10.64898/2026.08.28.747835

**Authors:** Maite Freire-Delgado, Diede de Haan, Stéphane Tachon, Anastasia D. Gazi, Mylan Ansel, Eva K. Pillai, Marvin Albert, Chantal Combredet, Audrey Salles, Jean-Yves Tinevez, Thibaut Brunet

## Abstract

Animal cells extensively remodel their cytoskeleton during differentiation and can notably switch between two major motility modes: flagellum-based swimming and actin-based crawling. We previously showed that choanoflagellates, the closest living relatives of animals and classically viewed as obligate flagellated swimmers, can retract their collar complex and adopt an amoeboid form within seconds under spatial confinement, independently of regulated gene expression. Here, using live imaging, ultrastructural expansion microscopy, and cryo-electron tomography in *Salpingoeca rosetta*, we identify rapid, cell-wide cytoskeletal remodeling as the ultrastructural basis of this switch. Unconfined choanoflagellates lack a detectable actin cortex but display an apical flagellum and cortical microtubules, with F-actin being largely restricted to microvilli. Confinement triggers calcium release from intracellular stores, which induces microvillar retraction and absorption of microvillar material into the cell body, including actin, ezrin-radixin-moesin 1, and plasma membrane. Remodeling of the internalized F-actin and repurposing of associated proteins supports *de novo* actin cortex formation, which is necessary for amoeboid motility. In parallel, cortical microtubules are disassembled, and the reabsorbed microvillar plasma membrane increases the surface area of the cell body, allowing the cell to flatten under confinement. Cryo-electron tomography reveals stepwise actin reorganization from internalized microvillar bundles to a cortical contractile meshwork combining bundles and scattered filaments. This work reveals considerable ultrastructural plasticity in the cytoskeletal architecture of choanoflagellates and supports an ancestral role for microvilli as reservoirs of membrane and cytoskeleton to potentiate cell phenotypic transitions.

## Introduction

Cytoskeletal architecture and activity underlie nearly every aspect of animal organismal form and function, from morphogenesis to cell differentiation. [1–5]. Moreover, a multicellular organism is a complex environment in itself, presenting cells with a vast range of forces and stimuli to navigate. To perform these functions, animal cells alternate between diverse phenotypes and modes of motility, including epithelial, mesenchymal, amoeboid, and flagellate cells, each with its characteristic cytoskeletal organization [6,7]. Interestingly, this cytoskeletal plasticity likely predated animals in evolution: microbial holozoans, the closest relatives of animals, can often alternate between diverse phenotypes and modes of motility in their life history [8–18].

Two of the most conserved modes of motility are flagellar swimming and F-actin-mediated cell crawling, which are broadly distributed in eukaryotes and conserved in animals [13,19]. This has led to the hypothesis that the unicellular precursor of animals might have been an amoeboflagellate alternating between (or combining) these two modes of motility [20,21], which might have resembled the amoeba *Naegleria* [22] or certain close relatives of metazoans with amoeboflagellate characteristics (such as some filastereans and ichthyosporeans [16,23–25]). In support of this model, we recently found that choanoflagellates, the closest relatives of animals, despite having long been seen as obligate flagellated swimmers, could switch to an amoeboid form when subjected to spatial confinement: they retract their flagellum, activate myosin II contractility, and engage in blebbing-based cell deformation [11]. This switch is conserved across most (but not all) choanoflagellate diversity and enables escape from confined microenvironments [11].

How this amoeboid switch is triggered remains an open question. Once exposed to confinement, cells begin blebbing in less than a minute, quickly followed by flagellar retraction after a few minutes. This process appears far too fast to depend on regulated gene expression – and indeed, inhibitors of transcription and translation fail to stop the transition, pointing toward a post-transcriptional and post-translational mechanism [11].

Here, we show that the amoeboid switch relies on a rapid and global reorganization of the choanoflagellate cytoskeleton (Fig. 1A). Upon confinement, calcium is released from intracellular stores, setting in motion a signaling cascade that promotes microvillar disassembly and releases microvillar components that serve as building blocks for the construction of the amoeboid architecture. First, the cortical microtubule cage detaches from the plasma membrane, microvilli retract, and their content – including actin, actin-associated proteins, and plasma membrane – is internalized and flows into the cell body. Second, the formerly microvillar actin undergoes stepwise remodeling, which we visualize by cryo-ET: reabsorbed microvillar actin bundles are progressively converted into a meshwork combining bundles and individual filaments, thus resulting in a canonical, contractile F-actin cortex underneath the plasma membrane. Meanwhile, the flagellar and cortical microtubules depolymerize, leaving only the microtubule-organizing center (the former flagellar basal body) intact. Overall, this supports an unexpected – and possibly pre-metazoan – role for microvilli as potentiators of phenotypic plasticity.

**Figure 1.**
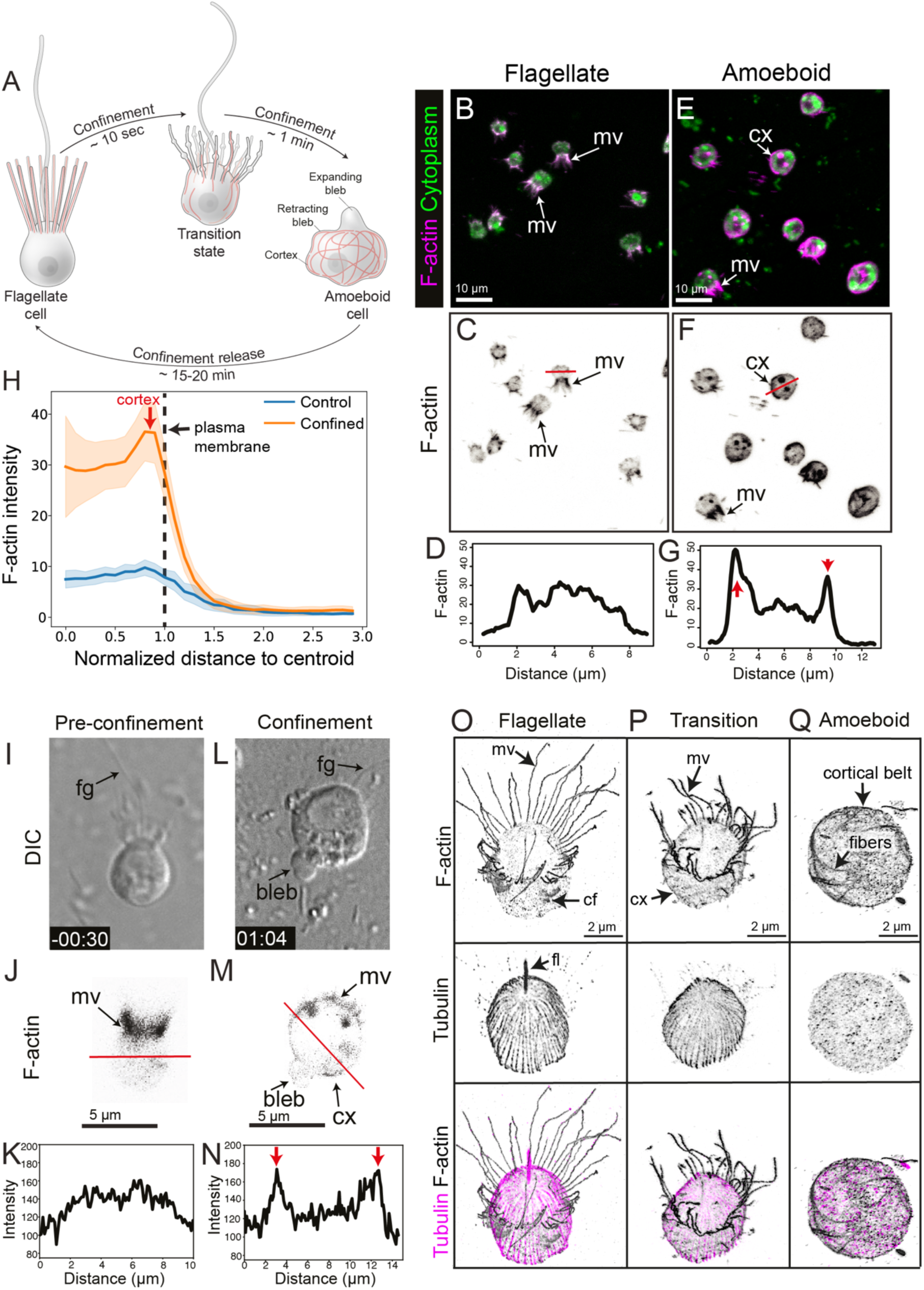
*S. rosetta* forms an actin cortex *de novo* under confinement. (A) Confinement induces global reorganization of the F-actin cytoskeleton. In unconfined flagellates, F-actin is almost wholly restricted to microvilli. Confinement induces flagellar and microvillar retraction, alongside internalization of microvillar F-actin bundles into the cell body. Actin remodeling results in the formation of a contractile F-actin cortex that supports blebbing-mediated cell deformation. (B-H) One-step fluorescent staining of *S. rosetta* reveals that F-actin is restricted to microvilli in flagellate cells, but forms a cortex under confinement. (B,C,E,F): maximum Z-projections of confocal stacks. (D,G): Linescans displaying F-actin staining intensity through the cell bodies of example cells across the red lines in C and F (respectively). F-actin was stained with Alexa Fluor 488-phalloidin and cytoplasm with FM 1-43 FX (a membrane dye that redistributes to the cell body upon fixation). (H): Quantification of F-actin distribution in the cell body of flagellate and amoeboid cells. Raw F-actin intensity (per cell) is plotted as a function of normalized distance to the cell centroid (1 corresponding to the cell membrane). N=54 control cells and N=36 confined cells. (I-N) Live imaging of an *S. rosetta* cell expressing the fluorescent F-actin marker LifeAct-mCherry reveals F-actin in microvilli prior to confinement and in the cortex after confinement. Note the presence of short residual microvilli in in the amoeboid cell. Time: min:sec. Linescans in (K,N) are along the red lines in (J,M) respectively. (O-Q) Ultrastructural expansion microscopy reveals cytoskeletal remodeling. (O) Flagellates display cortical microtubules, a tubulin-filled apical flagellum, F-actin-filled apical microvilli, and cytoplasmic actin foci (which might be endocytotic patches). (P) A transition-state cell lacking a flagellum and showing partly retracted microvilli, an incipient F-actin cortex, and cortical microtubules. (Q) A fully amoeboid cell showing an extensive F-actin cortex forming distinct fibers, notably as a circumferential ‘belt’, but also in the rest of the cell. No microtubules or microvilli are visible. In linescans (G,N): red arrows indicate cortex. In all panels: mv: microvilli; fg: flagellum; cf: cytoplasmic foci; cx: cortex.

## Results

### *S. rosetta* forms an actomyosin cortex *de novo* under confinement

To investigate the cytoskeletal architecture of the choanoflagellate cell (and notably of the collar complex, which is often disrupted by washing steps), we developed a one-step fixation/staining protocol for F-actin and cell body in *S. rosetta*. We observed F-actin in the microvillous collar, occasional filopodia, and occasional puncta in the cell body (possibly representing endocytic patches [26]), but were surprised to not see an F-actin cortex underneath the plasma membrane (Fig. 1B-D) – although such a cortex is a common feature of animal cells and of diverse unicellular amoebae [27,28]. This absence was surprising given the ability of *S. rosetta* to bleb under confinement [11], an activity that usually requires a contractile actin cortex [29,30]. We thus repeated one-step staining on choanoflagellates that had been confined for a few seconds in a 2 µm space using microbeads and, this time, noticed an unmistakable F-actin cortex (alongside retracted and scattered microvilli, as previously reported; Fig. 1E-G). Quantitative image analysis confirmed the presence of a peak of F-actin staining intensity at the periphery of amoeboid, but not flagellate, cells (Fig. 1H).

This observation suggested that *S. rosetta* might have the ability to form an F-actin cortex “on demand” under confinement. To independently assess this conclusion, we performed live imaging of transgenic *S. rosetta* expressing the live F-actin marker LifeAct-mCherry (Supp. Movie 1). Consistent with observations of fixed samples, we could not detect an F-actin cortex prior to confinement (Fig. 1I-K) but saw a cortex form within a few seconds of confinement and persist in actively blebbing cells (Fig. 1L-N).

Finally, we investigated the cytoskeletal ultrastructure of *S. rosetta* by superresolution (structured illumination microscopy, or SIM; Fig. S1) and by ultrastructural expansion microscopy (U-ExM; Fig. 1O-Q), taking advantage of a recently developed F-actin probe for expansion [31]. Flagellate cells were marked by a cage of cortical microtubules, microvillar F-actin, and a tubulin-filled apical flagellum (Fig. 1O, Fig. S1). F-actin puncta were once again observed in the cell body, but an actin cortex remained undetectable in both SIM and U-ExM (Fig. 1O, Fig. S1). U-ExM of the transition state revealed a retracted flagellum, partly retracted microvilli, and an incipient actin cortex (Fig. 1P). In fully amoeboid cells observed in U-ExM, microvilli were absent, but a conspicuous actin cortex was visible as thick fibers of F-actin underneath the plasma membrane (Fig. 1Q). Some actin fibers formed a peripheral “belt” within the confinement plane while others, of variable orientations, formed a loose network underneath other parts of the plasma membrane. Interestingly, while cortical microtubules coexisted with the F-actin cortex in the transition state (as previously reported [11]), fully amoeboid cells lacked observable microtubules (Fig. 1Q).

### Cells plated on high poly-D-lysine concentrations form an actin cortex

Taken together, these observations suggest that an F-actin cortex forms *de novo* under confinement during the flagellate-to-amoeboid switch in choanoflagellates. This stands in contrast to our own earlier assumption that the cortex pre-existed in the flagellate form [11] and motivated us to re-assess the evidence that had led to our earlier model. A key difference with the present study is that our earlier study involved cells attached to the substrate with a higher concentration of poly-D-lysine than in the experiments above (see Material and Methods). We hypothesized that this stronger poly-D-lysine-mediated adhesion might have unwittingly flattened the cells onto the coverslip, deforming them enough to cause cortex formation. To test this, we plated cells expressing LifeAct-mCherry on wells coated with a range of poly-D-lysine concentrations. While cells lacked an F-actin cortex at the lowest poly-D-lysine concentrations (Fig. S2A-D), intermediate concentrations induced formation of a conspicuous cortex coexisting with (shortened) microvilli (Fig. S2E-G). The highest concentrations induced a fully amoeboid phenotype, complete with F-actin cortex, microvillar retraction, and blebbing (Fig. S2H-K). We conclude that the F-actin cortex observed in our prior study had been induced by the method used for cell immobilization and did not actually pre-exist in freely swimming flagellates.

### Confinement-induced calcium signaling is necessary and sufficient for the amoeboid switch

A key question concerns the transduction pathway by which *S. rosetta* senses confinement to activate the amoeboid switch. In animal cells [32,33] and amoebae [27], cell confinement is often transduced (at least in part) by calcium signaling. While earlier experiments had failed to uncover clear evidence for calcium signaling in the amoeboid switch [11], we took advantage of the recent establishment of a transgenic strain of *S. rosetta* expressing the fluorescent calcium reporter RGECO [34] to revisit this question.

In the absence of confinement, cells displayed spontaneous and occasional calcium pulses (Fig. 2A-B), consistent with a previous study that reported such pulses and showed that they resulted in brief cessation of flagellar beating [34]. Spontaneous pulses were asynchronous between cells. By contrast, confinement caused almost all cells to undergo one to three calcium pulses within 10 to 15 seconds (Fig. 2C-E; Supp. Movie 3). Compared to spontaneous pulses, confinement-induced calcium pulses were longer (∼2-3 seconds, rather than <1 second) and more intense (ΔF/F_0_∼0.2, rather than 0.05 to 0.1). To test whether confinement-calcium signaling was causally linked to the amoeboid switch (rather than merely correlated), we artificially forced calcium influx using ionomycin. This caused a strong and synchronous calcium pulse (Fig. 2F-G; Supp. Movie 4) but also, remarkably, recapitulated all features of the amoeboid switch: ionomycin-treated cells retracted their collar and underwent blebbing within seconds (Fig. 2H-I; Supp. Movie 5). Furthermore, observation of ionomycin-treated cells expressing LifeAct-mCherry revealed formation of an actin cortex immediately after microvillar retraction (Fig. 2J; Supp. Movie 6). Finally, ionomycin-treated cells expressing a fluorescently tagged myosin light chain (MRLC-mTFP) showed condensation of myosin II into cortex-associated fibers and foci (Fig. S5A-F; Supp. Movie 7), a previously described hallmark of the amoeboid switch [11].

**Figure 2.**
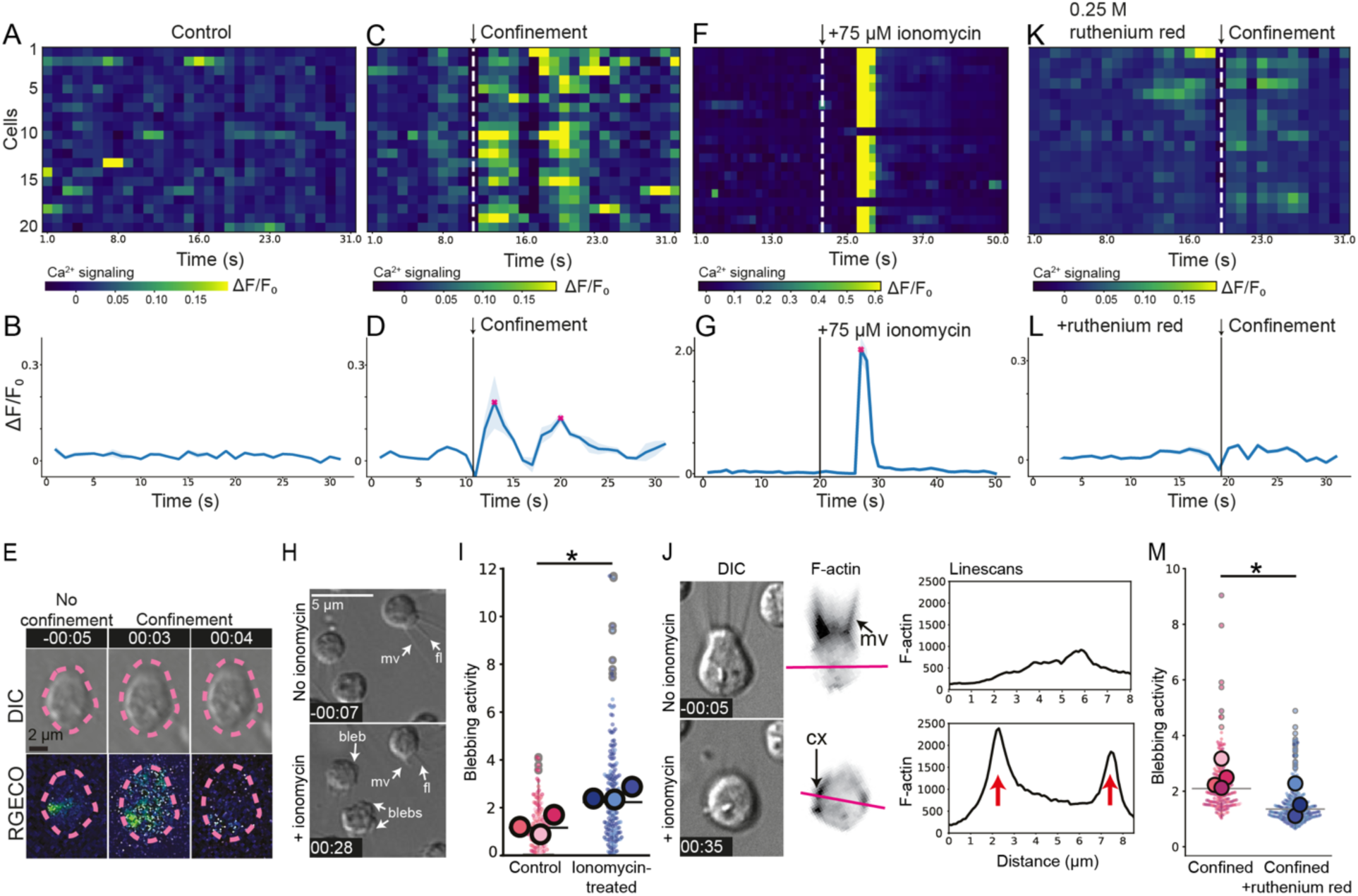
Confinement-induced calcium signaling is necessary and sufficient for the amoeboid switch. Across panels A to L, calcium signaling is visualized using the genetically encoded fluorescent calcium sensor RGECO [34]. (A, C, F, K) are heatmaps encoding time-course calcium signaling activity per cell (ΔF/F_0_ of RGECO fluorescence; see Methods) and (B, D, G, L) show averaged activity across all cells. Ribbon: standard deviation, red cross: peak. (A-B) Unconfined cells undergo spontaneous, asynchronous calcium pulses, as previously reported [34]. (C-D) Confinement is followed by one to three intense and sustained calcium pulses across almost all cells. (E) Micrographs of RGECO signal in a cell before, during and after confinement. Color-code is the same as in panel C. (F-G) Addition of ionomycin into the culture medium results in a strong, synchronous calcium pulse within 5 seconds (presumably reflecting the time needed for the compound to diffuse to the cells). Note that the color scale of the heatmap, and the y-axis scale of the graph, are different than in other panels due to the high intensity of the ionomycin-induced pulse. (H-J) Timelapse microscopy (H,J) reveals that ionomycin addition induces blebbing (H,I), microvillar retraction and cortex formation (I,J). (I) *p*=1.5% by Student’s *t*-test. (K-M) Treatment with the calcium signaling inhibitor ruthenium red almost completely abolishes spontaneous and confinement-induced calcium pulses (K,L) as well as blebbing under confinement (M). *p*=3.5% by Student’s *t*-test. (I,K) are SuperPlots where large dots are averages of individual experiments (biological replicates) and small dots are individual cells (technical replicates).

We then set out to test whether calcium signaling was necessary for the amoeboid switch. In an earlier study, we had failed to inhibit the amoeboid switch with the cell-permeant calcium chelator BAPTA-AM combined with EGTA to deplete both extracellular and intracellular calcium [11]. We reproduced that experiment using the RGECO strain to monitor calcium signaling and found that, unexpectedly, the combined treatment did not abolish spontaneous (Fig. S5G-H) nor confinement-induced (Fig. S5I-J) calcium pulses. One possible explanation might be sequestration of BAPTA-AM in the food vacuole (away from the cytoplasm): indeed, a fluorescent BAPTA-AM analog (BAPTA-1 Oregon Green 488 AM) accumulated in the vacuole but was undetectable in the cytoplasm (Fig. S5K). This suggests that previously attempted treatments had not efficiently inhibited calcium signaling.

Using the RGECO strain to screen for more efficient inhibitors, we found that high concentrations of the broad-spectrum calcium channel inhibitor ruthenium red [35] strongly reduced both spontaneous and confinement-induced calcium pulses (Fig. 2K-L). Ruthenium red treatment also nearly abolished blebbing under confinement (Fig. 2M; Supp. Movie 8; Supp. Movie 9), supporting the idea that calcium signaling is both necessary and sufficient for the amoeboid switch.

### Cortex formation occurs by repurposing of microvillar cytoskeleton

*De novo* cortex formation in *S. rosetta* raises the question of the origin of cortical actin. Since choanoflagellates retract their microvilli, their most prominent actin-based structure, during the amoeboid switch, one straightforward hypothesis is that microvillar actin (and actin-associated proteins) might be repurposed to form the cortex. Indeed, timelapse imaging of cells expressing LifeAct-mCherry before and during confinement reveals that microvillar F-actin appears to invade the apical pole of the cell body, following the outline of the plasma membrane and apparently continuously transitioning into an F-actin cortex (Fig. 3A; Supp. Movie 2). Quantification of F-actin intensity confirmed that F-actin localization is displaced from the microvilli to the cell body after confinement (Fig. 3B). Close inspection of U-ExM micrographs showed that F-actin at the base of retracting microvilli lay directly underneath the plasma membrane of the cell body, suggesting direct conversion of microvillar actin into cortical actin (Fig. 3C, Fig. S3).

**Figure 3.**
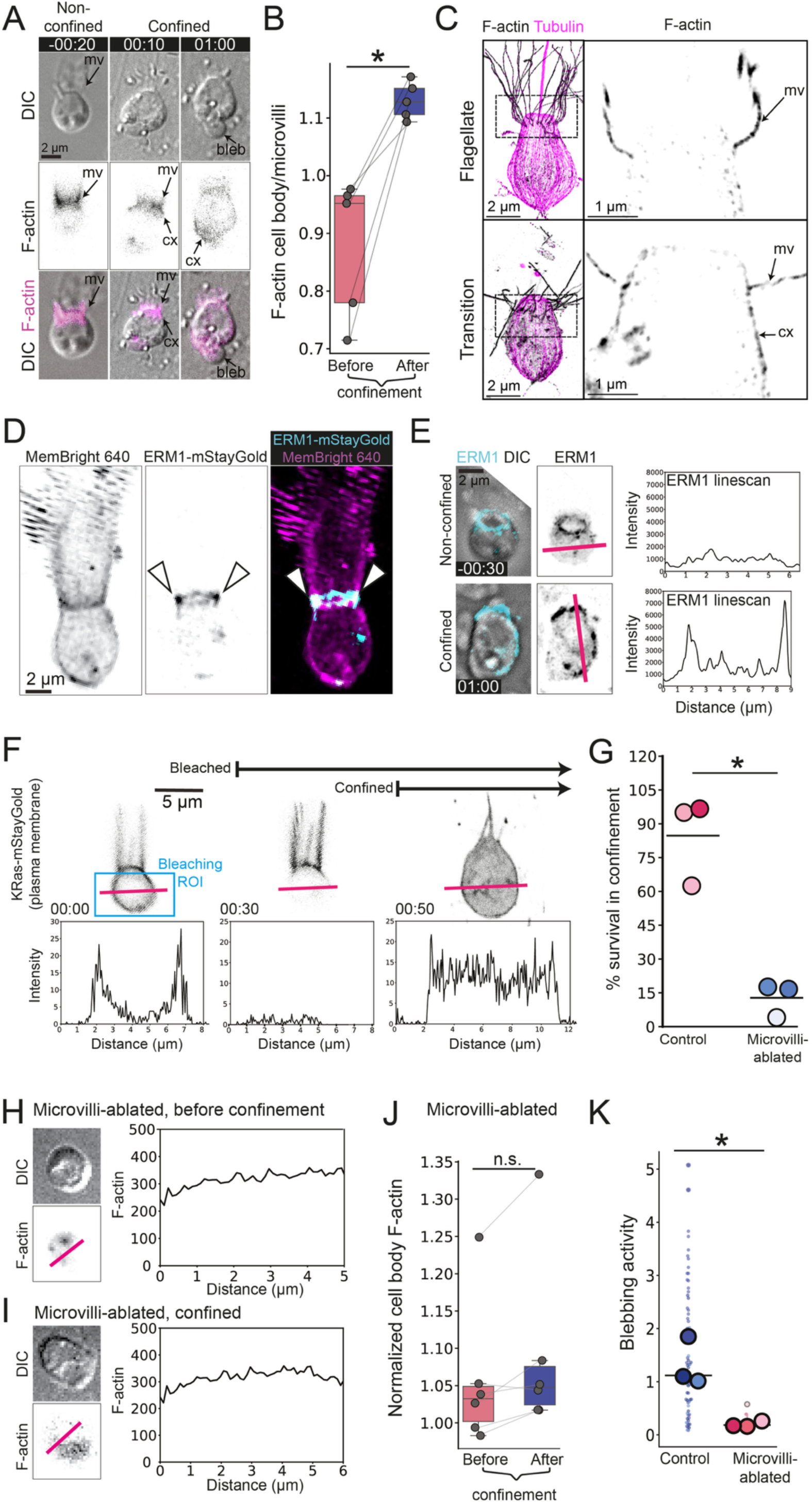
Microvillar actin, actin-associated proteins, and plasma membrane are reabsorbed and repurposed during the amoeboid switch. (A) Stills from a timelapse movie of an *S. rosetta* cell before and after confinement, showing internalization of microvillar actin before formation of an F-actin cortex across the whole cell body. Note the apparent continuity at 00:10 between the retracting microvillus and the forming apical cortex. The cell expresses LifeAct-mCherry. (B) Quantification of the cell body/microvilli ratio of LifeAct-mCherry intensity reflects the switch from a microvillus-dominated to a cortex-dominated actin cytoskeleton. Each dot represents an individual cell. *p*=0.8% by a paired Mann-Whitney test. (C) Ultrastructural expansion microscopy shows that F-actin in the microvilli, but not in the cortex, in flagellate cells. In transition cells, microvillar F-actin is reabsorbed in close contact with the plasma membrane, where the cortex will form. See also Fig. S3. (D) ERM1 is specifically located in the proximal section of microvilli in non-confined flagellate cells. ERM1 is visualized by endogenous tagging (ERM1-mStayGold) and cell outline by the vital dye MemBright 640. (E) ERM1 relocates from microvilli to the cortex upon confinement. Right: linescan across the red line immediately on the left. (F) Microvilli serve as a reservoir of plasma membrane that is reassigned to the cell body under confinement. The cell body of a cell expressing the plasma membrane marker KRas-mStayGold is bleached prior to confinement (blue box in 00:00; ROI: region of interest), leaving only the microvillar (and flagellar) plasma membrane fluorescent. Fluorescence in the cell body reappears immediately upon confinement, indicating that the plasma membrane of reabsorbed microvilli has been reassigned to the cell body. (G) Microvilli-ablated cells show compromised survival under confinement. *p*=1.4% by by Student’s *t*-test. Each dot represents an independent experiment. (H-I) Microvilli-ablated cells that survive confinement fail to form an F-actin cortex. Panels on the right: F-actin line scan across red line in the panels on the left. (J) No F-actin cortex forms in the cell body of surviving microvilli-ablated cells. The plot depicts LifeAct-mCherry intensity in the cell body normalized by extracellular noise. Compare to non-ablated controls in B. Each dot represents an individual cell. (K) Microvilli-ablated cells that survive confinement do not bleb. SuperPlot [75] where large dots represent average values of each biological replicate and small dots represent individual cells. *p=*5.0% by Student’s *t*-test.

To directly test the repurposing hypothesis, we initially attempted to use photoconversion to track microvillar actin before and after confinement. We reasoned that photoconverting an F-actin marker such as LifeAct would not necessarily be suitable to track actin itself, as such markers can dynamically bind and unbind microfilaments [36]. Therefore, we set out to directly tag *S. rosetta* actin with the photoconvertible fluorescent protein kikGR [37]. However, while we managed to endogenously tag one of the three *S. rosetta* endogenous actin loci (PTSG_05705) and while the resulting kikGR-actin fusion protein was expressed and photoconvertible, it virtually did not incorporate in microvilli (perhaps due to inefficient polymerization), precluding the tracking experiment (Fig. S4).

As a proxy to the localization of actin itself, we tracked actin-associated proteins during the amoeboid switch. In animal cells, certain actin-associated proteins play a dual role in both microvilli and cortex: this is the case of the actin-membrane linkers of the ezrin-radixin-moesin (ERM) family [38,39]. To investigate ERM localization in *S. rosetta*, we adapting a selection-based method for gene editing [40–42] to endogenously tag the ERM protein PTSG_04779 (hereafter ‘ERM1’) at its C terminus with the fluorescent protein mStayGold. ERM1 was restricted to the base of microvilli in flagellate cells (Fig. 3D) but broadly redistributed to the cortex upon confinement (Fig. 3E). Thus, cortex formation involves repurposing of microvillar actin-associated proteins.

### Reabsorbed microvillar plasma membrane increases the surface of the cell body

Beyond F-actin, microvilli are also covered in plasma membrane, which we reasoned might also be reassigned to the cell body under confinement. Indeed, confinement should be expected to create a need for additional plasma membrane in the cell body: while unconfined choanoflagellates are ovoid, confined flagellates have a flattened shape with a higher surface-to-volume ratio. Importantly, published electron microscopic observations [43–45] suggest that the plasma membrane of choanoflagellates lacks folded micro-structures such as the caveolae of animal cells, which can unfold under stretch and serve as a membrane reservoir [46]; moreover, one key protein for caveolae formation, caveolin, is animal-specific [47]. We set out to test the idea that microvillar retraction might meet (at least part of) the demand for extra plasma membrane under confinement. We photobleached the cell body of a strain of *S. rosetta* expressing the plasma membrane marker KRas-mStayGold [48], thus restricting fluorescent membrane to the collar (Fig. 3F). After confinement, fluorescence immediately redistributed to the cell body (Fig. 3F), indicating that reabsorption of microvillar membrane contributed to increasing cell body surface area.

### Microvilli are necessary for cortex formation and promote survival under confinement

The data above suggest that microvilli play an unexpectedly central role in the amoeboid switch, which notably involves general repurposing and redistribution of microvillar material in the cell body (including membrane, actin-associated proteins, and likely actin itself). We set out to test how microvillar ablation would affect the switch. We ablated the microvilli of LifeAct-mCherry-expressing cells by papain digestion and confined the resulting demicrovillated cells, notably monitoring the appearance (or lack thereof) of a cortex. The first and most conspicuous effect was a dramatic reduction of survival under confinement: while ∼80% of control cells survived, only ∼15% of ablated cells did (Fig. 3G). We scrutinized surviving cells and noticed that they had failed to assemble an actin cortex (Fig. 3H-J) and did not bleb (Fig. 3K), despite visible cell flattening confirming that confinement had taken place. This supports the idea that repurposing of microvillar material is crucial for cortex formation and cell survival under confinement. The effect on survival might reflect both a contribution of the cortex itself to withstanding compressive forces and/or the need for additional plasma membrane to support cell flattening and prevent passive bursting.

### Blotting before plunge-freezing induces cortex formation, but high-pressure-freezing preserves the native flagellate ultrastructure

To further test the idea that microvillar actin might be remodelled to form the amoeboid cortex, we set out to investigate the fine structure of the cytoskeleton before, during and after the amoeboid switch, by cellular cryo-electron tomography (cryo-ET [49]). To induce the amoeboid switch, we treated LifeAct-mCherry-expressing cells for 1 minute with 75 µM ionomycin and vitrified them by plunge-freezing (Fig. 4A). We also plunge-froze untreated control cells in parallel, and monitored general actin architecture of both types of cells by cryo-confocal microscopy prior to sectioning. Surprisingly, although we expected untreated cells to display a native flagellate phenotype, an F-actin cortex was visible and microvilli often seemed to be in the process of retracting, suggesting that untreated cells had initiated an amoeboid transition before being fixed (Fig. 4B-C). In hindsight, we identified the likely cause of this switch as the 9-second blotting step that had preceded plunge-freezing, which had confined cells in a thin liquid film on the EM grid. Although necessary to minimize volume and achieve vitrification, this step appeared to have sufficiently confined cells to make them amoeboid (consistent with our earlier findings on choanoflagellates trapped in a thin liquid film [11]). Ionomycin-treated samples had a more extensive amoeboid phenotype (without observable microvilli, a marked F-actin cortex and frequent blebs; Fig. 4D-E), consistent with the blotting step having only lasted 9 seconds while ionomycin treatment had lasted 1 minute.

**Figure 4.**
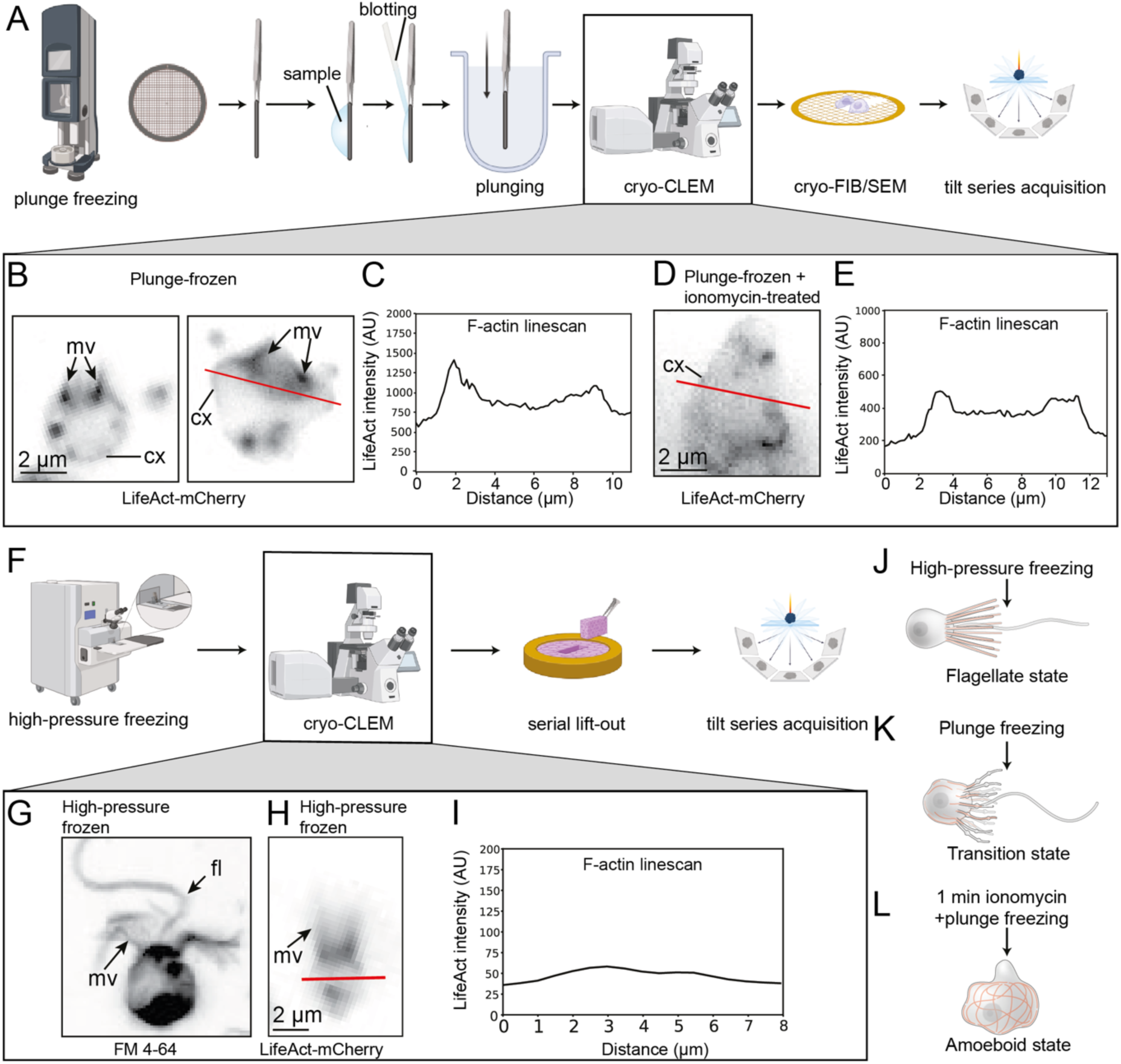
Plunge-freezing induces cortex formation and high-pressure-freezing preserves the native flagellate ultrastructure. (A) Plunge-freezing workflow. (B) Plunge-frozen cells display an F-actin cortex and microvilli at various degrees of reabsorption, corresponding to the early “transition” stage of the amoeboid switch (compare Fig. 1P). (C) F-actin intensity line scan across the red line in B. (D) Plunge-frozen cells that have been previously treated for one minute with ionomycin display an F-actin cortex and lack microvilli, corresponding to a more advanced stage of the amoeboid switch (compare Fig. 1Q). (E) F-actin line scan across the red line in D. (F) High-pressure freezing workflow. (G) High-pressure freezing preserves collar and flagellum morphology. (H) High-pressure frozen cells display well-preserved microvilli and lack an F-actin cortex. (I) F-actin line scan across the red line in H. (J-L) Distinct cryofixation procedures preserve the flagellate (J), transition (K) and amoeboid (L) phenotypes. Cartoons of equipment in A and F were adapted from BioRender.com. In all panels: mv: microvilli; cx: cortex.

To better preserve the native flagellate architecture, we turned to another cryo-fixation method, high-pressure freezing (Fig. 4F). In this approach, cells are vitrified at high pressure in planchettes containing a sufficient liquid volume to avoid confinement. Cryo-confocal observation of high-pressure-frozen cells stained with the membrane marker FM-4 64 revealed maintenance of the collar, flagellum, and typical ovoid shape of flagellate cells (Fig. 4G). Observation of LifeAct-mCherry confirmed that F-actin was restricted to microvilli and cell body puncta, and that no observable cortex had formed (Fig. 4H-I). This came in further support of the idea that the amoeboid switch in our earlier assays had been induced by confinement before plunge-freezing and gave us a method for cryo-ET of unconfined flagellate cells. Guided by the fluorescent signal observed in cryo-confocal microscopy, we prepared lamellae of areas of interest in flagellates (by serial lift-out [50] or by the waffle method [51]) (Fig. 4J) and transitional/amoeboid cells (by cryo-focused ion beam (cryo-FIB) milling) (Fig. 4K-L).

### Cryo-ET reveals stepwise remodeling of actin nanoarchitecture

Flagellate tomograms revealed that microvilli contained abundant and tightly bundled F-actin (Fig. 5A, Supp. Movie 10). By contrast, in the cell body, cortical microtubules were readily observable underneath the plasma membrane, but no cortical actin microfilaments were detected (Fig. 5B, Supp. Movie 11). In early amoeboid cells, retracting microvilli appeared disorganized (Fig. 5C, Supp. Movie 12)) and F-actin displayed an intermediate architecture: microvillar bundles of microfilaments invaded the cell body, coming to lie underneath the plasma membrane (Fig. 5C-D, Supp. Movie 13). Between bundles, a few isolated actin microfilaments lay in diverse orientations (Fig. 5D). Formerly cortical microtubules were detached from the plasma membrane, below the forming cortex, and appeared less regularly organized than in flagellates (Fig. 5D). Finally, in fully amoeboid cells, microtubules were nearly absent from tomograms, with the notable exception of the centriole that serves as a basal body/MTOC (and a few surrounding microtubules; Fig. 5F). The cell body displayed an extensive cortex of F-actin, combining small bundles interspersed with more isolated microfilaments (Fig. 5E, Supp. Movie 14). Overall, these observations are in good agreement with U-ExM data (Fig. 1O, Fig. 3C). This is also consistent with our earlier finding that flagella regrow close to their initial position after confinement release [11], which had been interpreted as reflecting likely maintenance of MTOC presence and localization in amoeboid cells.

**Figure 5.**
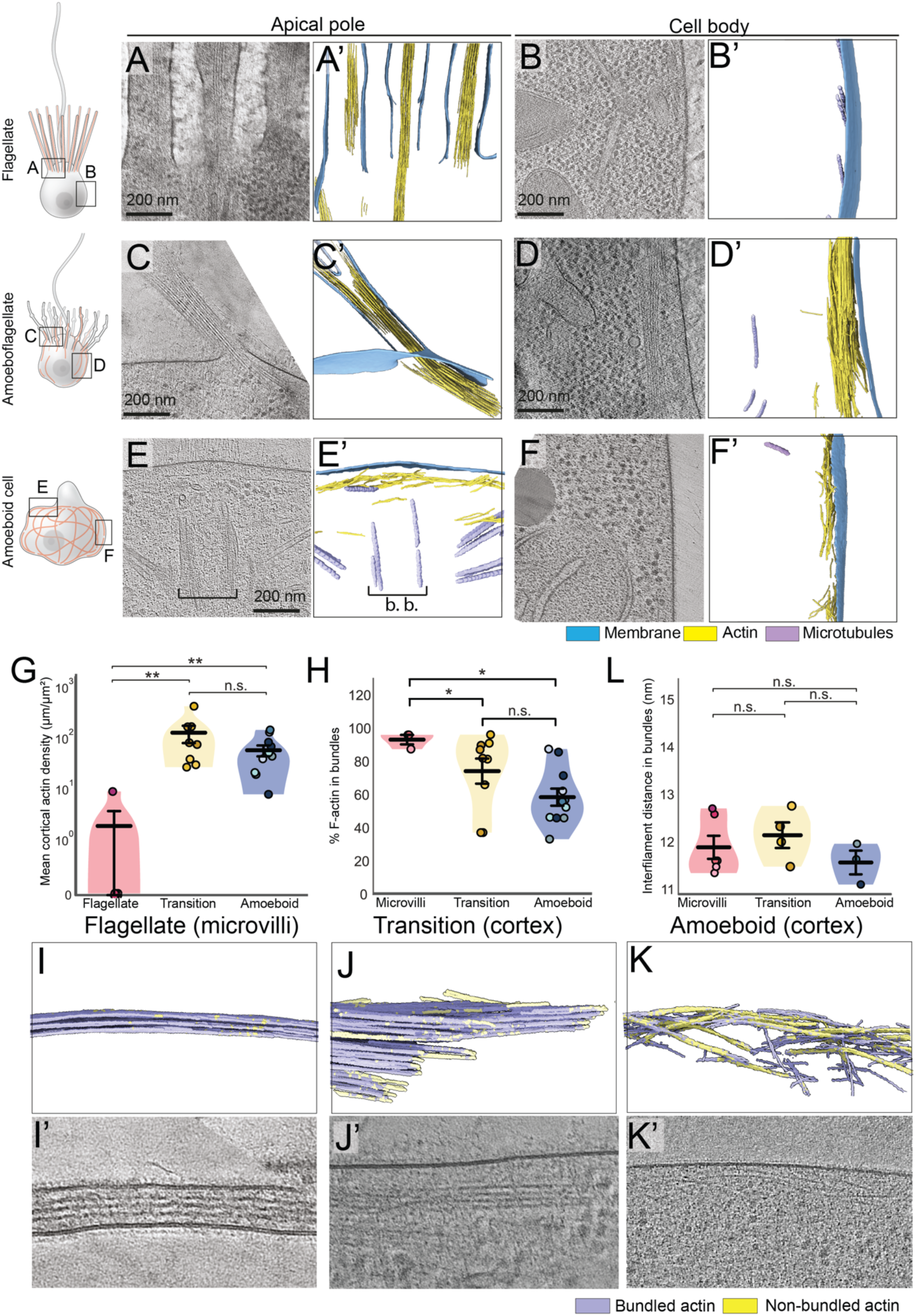
Cryo-ET reveals stepwise remodeling of actin nano-architecture. (A-F) Slices through reconstructed cryo-tomograms and segmented reconstructions (A’-F’) of cells in flagellate (A-B), transitional (C-D), and ameoboid (E-F) states. The apical pole initially bears microvilli characterized by tight bundles of parallel actin microfilaments (A-A’), which are reabsorbed and fragmented in the transition state (C-C’) to be replaced by an actin cortex (D-D’) in fully amoeboid cells, whose apical pole can only be recognized by the persistence of the former flagellar basal body (bracket). (E-E’). The cell body is initially devoid of detectable actin (B-B’) but comprises cortical microtubules underneath the plasma membrane. In transitional cells, the microtubules are detached from the plasma membrane (D-D’) and bundles of microfilaments, reminiscent of microvillar actin architecture, come to lie in close contact with the plasma membrane instead, forming an incipient cortex. (F-F’) A full-fledged F-actin cortex is visible in the cell body of amoeboid cells, with the former cortical microtubules having become nearly undetectable in tomograms, consistent with UExM results (see Fig. 1P). (G) Quantification of F-actin cortical density shows near-absence of cortical F-actin in flagellate cells but abundant microfilaments in transition and amoeboid states. Note that transition cells display more F-actin than fully amoeboid ones, suggesting that reabsorbed microvillar actin is sufficient to account for the whole pool of cortical actin found in amoeboid cells. *p*=0.4% and 0.3% for the flagellate/transition and flagellate/amoeboid comparisons respectively. (H-K) Fraction of bundled F-actin decreased continuously from the microvilli of flagellated cells (I-I’) to the cortex of transitional cells (J-J’) to the cortex of amoeboid cells (K-K’), consistent with the final cortical architecture being a mixture of bundled and scattered microfilaments. (I, J, K) are segmented reconstructions and (I’, J’, K’) are slices through the corresponding reconstructed tomograms. *p=*1.9% for the microvilli/amoeboid comparison in (H). (L) Architecture of F-actin bundles does not significantly change from microvilli to the final cortex, with conserved inter-filament spacing (core-to-core distance between adjacent microfilaments) close to 12 nm.

Quantification of cortical F-actin density per surface unit of plasma membrane (Fig. S6) confirmed that cortical actin was nearly absent in flagellate cells, peaked in abundance in the transition phase (when most or all microvillar F-actin was internalized in the cell body), and remained high in fully amoeboid cells (Fig. 5G). Percentage of bundled actin continuously decreased during the amoeboid switch: while microvillar F-actin was nearly 100% bundled (Fig. 5H-I), bundling fraction was 80% in the transition state (Fig. 5H-J) and 60% in the cortex of fully amoeboid cells (similar to the cortex of mouse fibroblasts [52]; Fig. 5H-K). Interestingly, inter-filament spacing (∼12 nm, core-to-core distance) was similar in microvillar and cortical bundles (Fig. 5L), suggesting presence of a shared cross-linker. An interesting candidate is fimbrin/plastin, which imposes a 12-nm spacing between microfilaments [53], is conserved in choanoflagellates [39], and bundles both microvillar and cortical actin in animals [38,53–55].

### Discussion and conclusion

The present study reveals the structural basis of the flagellate to amoeboid transition in *S. rosetta*: confinement induces retraction of collar microvilli followed by *de novo* assembly of a contractile actomyosin cortex and loss of most cortical microtubules. These results extend and complement our previous finding that the amoeboid switch in *S. rosetta* occurs within seconds to minutes and is independent of regulated gene expression [11].

The first central finding of this study is that cortex formation relies on repurposing of microvillar material. Live imaging, papain-mediated microvillar ablation, ERM1 relocalization, and membrane photobleaching together support a model in which retracting microvilli supply actin, actin-membrane linkers, and plasma membrane to the cell body, where these components are reorganized into a cortical actomyosin system. Cryo-ET further reveals that this is not an abrupt replacement of one architecture by another, but a stepwise transformation from bundled microvillar actin to a cortical meshwork combining bundles and individual filaments. In this sense, microvilli are not simply lost during the amoeboid switch, but provide necessary material for a new cellular architecture.

This complements our recent finding that the flagellum similarly switches function under intermediate levels of confinement (∼3 µm, sufficient to prevent swimming but not to induce flagellar retraction): in this intermediate state, the flagellum straightens and adheres to the substrate, repurposing the molecular motors of intraflagellar motility to support gliding motility and propel the cell at ∼1 µm/s [18]. We hypothesize that in real confined microenvironments, flagellar gliding (that provides motor force) might cooperate with blebbing (that mediates cell body deformation and might help navigate obstacles, pores or constrictions). Thus, both flagellum and microvilli are repurposed to navigate confined spaces. More generally, this suggests that choanoflagellates possess a far richer behavioural repertoire than classically assumed [56]. In nature, these fast and on-demand cytoskeletal reorganisation might support exploration of diverse microhabitats, and notably of fine interstitial spaces and liquid films within the soil, that are known to house complex microbial communities [57], including choanoflagellates [58]. Testing these hypotheses in more realistic and complex microenvironments is a stimulating avenue we aim to explore in the future.

A second main conclusion is that calcium signaling is both necessary and sufficient for this transition. Confinement elicits calcium pulses that are stronger and longer than the spontaneous pulses observed in unconfined cells, and artificial calcium mobilization by ionomycin is sufficient to trigger collar retraction, cortex formation, myosin II reorganization, and blebbing. Conversely, inhibition by ruthenium red abolishes both calcium transients and amoeboid behavior. The fact that confinement-induced pulses induce blebbing, but not spontaneous pulses (that cause brief flagellar immobilization instead [34,59]), suggests that not all calcium signals are functionally equivalent. Calcium is one of the most versatile intracellular second messengers, and cells decode calcium signaling through amplitude, frequency, duration, and spatial compartmentalization. In animal cells, this versatility enables calcium to regulate a broad range of physiological processes, from fertilization to proliferation, migration, secretion, neuronal activity and apoptosis [60]. It is thus unsurprising that calcium signaling might control diverse processes in choanoflagellates.

This repurposing of microvillar actin has parallels in animal cells. The most similar process is probably “enterocyte restitution”: after injury of an epithelial enterocyte monolayer, brush-border microvilli are dismantled and their actin is repurposed to generate a lamellipodium that allows partial epithelial-to-mesenchymal transition, directional cell migration, and wound closure [61–64]. Functional studies have shown that the microvillar actin-binding protein villin plays a central role in enterocyte restitution. Under resting conditions, villin functions as a microvillar actin bundler, whereas calcium elevation converts it into an actin-severing and capping protein that drives brush-border disassembly and releases actin for lamellipodium formation [63,64]. Villin itself also relocalizes from the apical domain to the lamellipodium in the process [63,64]. Although the molecular effectors of actin remodeling remain to be identified in choanoflagellates, the overall logic appears similar: calcium-triggered dismantling of microvilli supports the rapid emergence of a new motile state. Interestingly, choanoflagellate genomes encode two villin homologs [39,65], that remain functionally uncharacterized in *S. rosetta*. Relocalization of ERM1 during the choanoflagellate amoeboid switch resembles epithelial-to-mesenchymal transition in mammary gland cells, during which moesin (one of the three mammalian ERM paralogs) relocalizes from microvilli to migratory protrusions (Haynes et al, 2011). More broadly, this highlights the plasticity of actin networks and supports a view of actin assemblies as dynamic and interconvertible rather than fixed entities [66]. It also fits with the emerging concept that different actin networks within the same cell can compete for shared pools of monomers [67].

Beyond providing F-actin, microvilli also serve as a source of plasma membrane during the *S. rosetta* amoeboid switch – which is likely important because confinement imposes a flattened cell geometry with an increased surface-to-volume ratio and because *S. rosetta* lacks observable caveolae-like membrane stores. In animal cells, microvilli similarly act as membrane reservoirs in multiple contexts, including cellularization in fly embryos [68] or to buffer membrane tension during spreading, mitosis, or osmotic stress [69,70].

These findings have implications for choanoflagellate biology and early animal evolution. Choanoflagellate microvilli are generally considered to mediate bacterial prey capture [39] and environmental sensing [39,59,71]. Our results indicate they play an additional and hitherto unsuspected role as reservoirs of cytoskeletal and membrane material to potentiate rapid phenotypic transitions that do not rely on regulated gene expression or protein synthesis. Given the similar function of animal epithelial microvilli as a readily mobilizable source of actin and membrane, the function of microvilli as potentiators of phenotypic transition may have preceded animal multicellularity in evolution. More broadly, our results support the hypothesis that the unicellular ancestor of animals was a complex, phenotypically plastic organism that possessed multiple actin networks [28,72] and could alternate between (or combine) flagellated and amoeboid phenotypes. Such amoeboflagellate phenotypes are indeed common across close relatives of animals, including choanoflagellates [11], filasterans [23–25], and ichthyosporeans [16] – but are also found in certain more distantly related eukaryotes such as *Naegleria* [19]. Such ancestral cellular phenotypic plasticity may have paved the way for the evolution of animal cell types by temporal-to-spatial transition [8,20,21,73,74].

## Methods

**Table S1.**
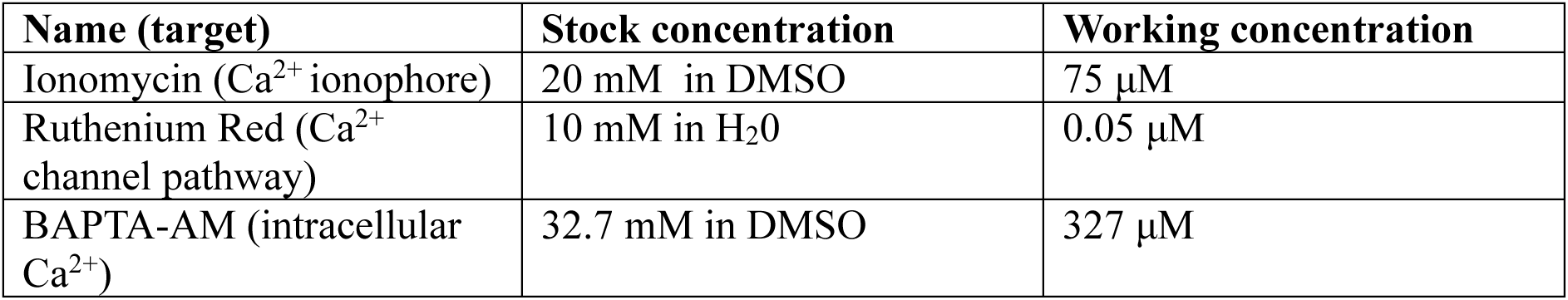
Drugs used in pharmacological assays.

Centrifugation steps were conducted in an Eppendorf 5415R microcentrifuge for small volumes (≤2 mL) and in a FisherBrand GT2R Expert centrifuge for large volumes (2–50 mL).

### Choanoflagellate culture

The chain/slow swimmer form of *Salpingoeca rosetta* [76] was cultivated in co-culture with its bacterial prey *Echinicola pacifica* (SrEpac; [77]) in 25 cm^2^ tissues culture flask containing 1X filter sterilized Artificial Seawater (ASW) prepared from Instant Ocean Powder (Aquarius System) supplemented with 5% SeaWater Complete Medium (SWC). Cultures were maintained in a Memmert IP410ecoplus incubator at 21°C. Humidity was controlled by placing a container with distilled water on the bottom shelf of the incubator. *S. rosetta* transgenic strains expressing LifeAct-mCherry [11], MRLC-mTFP [11], RGECO [34] and KRas-mStayGold [48] were all previously described and cryogenically stored in liquid nitrogen as previously described [40] using glycerol as a cryoprotectant [78].

### Preparation of *E. pacifica* Food Pellets

A 5 mL aliquot of an *Echinicola pacifica* culture in 100% SWC medium pre-grown from a glycerol stock was inoculated into 500 mL of 100% SWC medium. The culture was incubated for 2–3 days at room temperature on a rocking shaker until reaching an optical density (OD) of approximately 0.6. The bacterial suspension was then divided into 50 mL Falcon tubes and centrifuged at 4,000–5,000 × g for 30 minutes at 4 °C. The supernatant was carefully removed using a serological pipette to eliminate residual liquid. The bacterial pellet was weighed, and 1× ASW was added to achieve a final concentration of 20 mg/mL. The bacteria were resuspended by gentle pipetting, and 500 μL aliquots (corresponding to 10 mg of bacteria) were transferred into 1.5 mL Eppendorf tubes. These aliquots were centrifuged at 16,000 × g for 10 minutes at 4 °C, and the resulting supernatant was removed with a fine pipette tip. The bacterial food pellets were flash-frozen in liquid nitrogen and stored at –80°C.

### Dynamic Cell Confinement

To image *S. rosetta* cells under confinement, 5 mL of a dense culture (∼1e5 cells/mL) were pelleted by centrifugation at 2,000 × g for 15 minutes and resuspended in 1 mL of ASW. The concentrated suspension was placed in FluoroDishes (World Precision Instruments, #FD35-100) and covered with a coverslip covered in 2 µm micropillars connected to a suction line. The chamber was mounted in a one-well dynamic cell confiner (4Dcell, Montreuil, France) coupled to an Elveflow Vacuum/Pressure Generator and an Elveflow AF1 DUAL Vacuum/Pressure Controller. Confinement pressure was gradually decreased from −3 kPa to −10kPa with the vacuum/pressure controller. Confinement was released by gradually restoring pressure to -3kPa.

### Poly-D-lysine-mediated cell deformation

*S. rosetta* cells were plated on surfaces coated with different concentrations of poly-D-lysine (Poly-D-Lysine, Thermo Fisher Scientific, Cat. No. P0899) to generate different degrees of adhesion. The poly-D-lysine solution was prepared in distilled water by dilution of a 0.1 mg/mL stock and applied to the wells for 30 seconds before removal. No washing steps were added in this series of experiments. Cells were immediately plated onto the treated surfaces.

### Transmitted Light Live Imaging

Cells were imaged using differential interference contrast (DIC) microscopy with either a 40x or a 63x water immersion C-Apochromat 1.1 NA objective mounted on a Zeiss Observer Z.1 equipped with a Hamamatsu Orca Flash 4.0 V2 CMOS camera (C1140-22CU).

### Confocal Imaging

Samples were either imaged with a Leica Stellaris 5 confocal microscope equipped with an HC PL APO 63×/1.20 W CORR CS2 objective in Lightning super-resolution mode, or with a Plan-Apochromat 40X/1.31 Oil DIC (UV) VIS-IR M27 Zeiss objective using AiryScan MPLX SR-4Y and 2X line averaging1modes in an inverted Zeiss Axio Observer Z1/7 confocal microscope equipped with LSM9001Airyscan 2.

### Actin cortex quantification in confocal micrographs

To quantify cortical F-actin distribution, confocal z-stacks were optionally corrected for lateral drift in z and maximum projected. Cell bodies were segmented based on the membrane channel using Cellpose [79], including filtering of the segmentation labels using a size filter. Background intensities were estimated from the cell-free image regions and subtracted from the images. Actin intensities were then sampled radially from each cell center in rays sampled at an angular distance of 10 degrees. Distances from the cell center were normalized to the estimated cell radius and grouped in bins of width 0.1. Ribbons on the confined and unconfined intensity measurements represent half of the standard deviation across cells. Analysis was performed using custom Python scripts available under https://github.com/m-albert/choano_cortex_quantification [80].

### Structured Illumination Microscopy

Cells were fixed and stained as in [43], with the following modifications: coverslips (diameter 13 mm, Fisher Scientific 15893437) were not pre-coated with poly-L-lysine, and were manually coated instead with 50 mg/mL poly-D-lysine (applied for 30 seconds over the entire surface of the coverslip, removed, and washed three times with distilled water). Cells were stained using Alexa Fluor 488-phalloidin (10125092) and a mouse anti-tubulin antibody (Developmental Studies Hybridoma Bank, E7) detected with a highly cross-adsorbed Cy3 goat anti-mouse secondary antibody (InterChim XEQ110). After staining and washes, samples were mounted in ProLong gold antifade reagent (Fisher 11559306) and imaged on a Zeiss Elyra 7 microscope in the Photonic BioImaging facility of the Institut Pasteur.

### Bleaching

*S. rosetta* cells expressing the plasma membrane marker KRas-mStayGold [48] were seeded onto a 35 mm round dish (μ-dish 35 mm high, 81156, Ibidi) and imaged using a Leica Stellaris 5 confocal microscope. FRAP experiments were performed using the FRAP module in LAS X software. A region of interest (ROI) encompassing part of the cell body was selected for photobleaching. The ROI was bleached using a 405 nm laser (five iterations at maximal laser power).

### One-step staining of flagellate and amoeboid cells

One-step staining of flagellated *S. rosetta* cells was performed by concentrating the cells and fixing them in suspension. The following fixation solution was added: 4% paraformaldehyde (PFA; Electron Microscopy Sciences, 15710), 3.3× phosphate-buffered saline (PBS; Sigma-Aldrich, 806552), 10 mM EGTA, and 0.01% Triton X-100 (Thermo Fisher Scientific, A16046) prepared from a pre-diluted 10% (v/v) stock solution in distilled water. The fixation solution also included the following fluorescent dyes: (1) FM 1-43FX (Thermo Fisher Scientific, F35355), prepared as single-use aliquots (1 mg/ml) in water, stored at -20°C, and added at a 1:1,000 dilution. Alexa Fluor™ 488 Phalloidin (Invitrogen, A12379) was dissolved to 2 U/µL in DMSO, stored at -20°C, and added at a 1:300 dilution. Hoechst 33342 (Invitrogen H21486) was added at a 1:1,000 dilution.

One-step staining of confined cells was performed by mounting the cells between a small round coverslip (0.13–0.16 mm thickness; Fisher Scientific, 10513234) and a FluoroDish (35 × 23 mm; World Precision Instruments, FD35-100) with 1 µm microbeads as spacers (ThermoFisher C37274, diluted 1:100). Immediately after confinement, the edges of the round coverslip were secured to the large coverslip with a small amount of nail polish to maintain close apposition of the two coverslips and preserve cell confinement throughout the staining procedure. 10 µL of fixation solution was pipetted at the inlet side of the chamber to draw the solution by capillary action. Of note, a similar approach was published by an independent team [81] while we were working on this study, confirming the broad applicability of one-step staining.

### Ultrastructural Expansion Microscopy (U-ExM) – Cell fixation

The U-ExM protocol was adapted from [17,82,83]. 6 mm coverslips were placed in a 48-well plate and covered either with 30 µl of 10 % (0.01 mg/mL) poly-D-lysine for 30 s and washed once (flagellate condition) or with 60 % (0.06 mg/mL) poly-D-lysine for 30 s without being washed (transition and amoeboid conditions). 1 mL of *S. rosetta* culture (for each condition) was concentrated at 3,500 g for 4 min at room temperature and the supernatant was removed to keep the bottom 50 µL. The resulting concentrated culture was then deposited on the coverslip and incubated for 25 min to let cells settle and switch to the amoeboid state. Cells were then fixed with 1.7 % paraformaldehyde (Electron Microscopy Sciences 15714), 0.06 % glutaraldehyde (Electron Microscopy Sciences 16220) in 1X PHEM, 1.5X ASW buffer for 30 min before being washed twice with 4X PBS.

Buffers were prepared as follows: *(1)* PHEM buffer had been pre-emptively prepared as a 2X stock with the following recipe: 120 mM PIPE, 20 mM EGTA, 50 mM HEPES, 4 mM MgCl2, pH = 7.4. (*2)* PHEM-ASW buffer was prepared by diluting 2X PHEM 1:1 in 3X ASW (for 50mL: 25 mL 2X PHEM, 25 mL 3X ASW). *(3)* Finally, the fixation solution (1X PHEM 1.5X ASW) was prepared as follows: 2% paraformaldehyde, 0.075% glutaraldehyde in PHEM-ASW (for 2 mL: 666.7 µL 8% PFA (diluted in water) + 8 µL 25% GA in PHEM-ASW buffer).

### U-ExM – F-actin staining

F-actin was stained using the HAK-actin probe (Spirochrome SC019) with a protocol adapted from [31]. Cells were permeabilized for 10 min using 0.1% Triton X-100 in 4X PBS. Stock of prediluted HAK-actin solution at 1/100 in DMSO was freshly diluted at 1/10 in 4X PBS (1/1000 final) and a 15 µL drop for each coverslip was placed in parafilm. Coverslips were flipped on top of them and incubated for 1 h at room temperature before being replaced in the 48-well plate. Cells were then fixed and washed again as described above.

### U-ExM – Gelation

Cells were anchored in an acrylamide/paraformaldehyde solution (1% acrylamide, 0.7% paraformaldehyde) overnight at 37°C. Monomer solutions composed of 19% (wt/wt) sodium acrylate (Sigma-Aldrich 408220), 10% (wt/wt) acrylamide (Sigma-Aldrich, A4058), 0.1% (wt/wt) N,N′-methylenebisacrylamide (Sigma-Aldrich M1533) in PBS were used with 10% (wt/wt) Tetramethylethylendiamine (TEMED ThermoFisher 17919) and 10% (wt/wt) APS (ThermoFisher 17874) for gelation, and gels were allowed to polymerize for 1 h at 37°C in a moist chamber. For denaturation, gels were transferred to denaturation buffer (50 mM Tris, pH 9.0, 200 mM NaCl, 200 mM SDS) for 15 min at RT and then incubated at 95°C for 1 h. Following denaturation, expansion was performed with three water exchanges of 20 min each. After expansion, gel diameter was measured to determine the expansion factor, and scale bars have been rescaled according to the gel expansion factor to indicate actual size.

### U-ExM – Staining and imaging

Prior to staining, gels were first shrunk in 1X PBS for 15 min. Primary antibodies for tubulin (guinea pig, AA344 ABCD Antibodies) and HA (rabbit, Protein Tech 51064-2-AP) were diluted at 1/200 and 1/100 respectively in 3% PBS, 0.1% Tween 20 (PBST) with 3% BSA, and incubated overnight at 37°C, 80 rpm. Gels were then washed three times for 10 min, 80 rpm in PBST. Anti-guinea pig (Alexa Fluor 647 ABCAM ab150187-500 µg) and anti-rabbit (AlexaFluor 680 Invitrogen A21109) secondary antibodies were both diluted at 1/300 in PBST with 3% BSA, and incubated for 3 h at 37°C, 80 rpm in a dark moist chamber. Gels were then washed three times for 10 min, 80 rpm in PBST and stained overnight at 4°C, 80 rpm in dark with BODIPY (2mM stock in DMSO, ThermoFisher D7540) and NHS-Ester 405 (10 mg/mL stock in DMSO, DyLight™ 46401) diluted at 1/300 and 1/600 respectively in 1 X PBS (not shown in figure). Finally, gels were re-expanded with three washes of 15 min in water. Gels were mounted in FluoroDishes covered with 100% Poly-D-Lysine for 30 min and imaging was performed using Leica Stellaris 5 confocal microscope with an HC PL APO 63x/1.20 W CORR CS2 objective and Lightning super-resolution mode.

### Ionomycin Treatment

To increase intracellular Ca²⁺ levels and induce the amoeboid switch in the absence of confinement, ionomycin (ThermoFisher Scientific I24222) was added to cultures. A range of concentrations from 10 µM to 200 µM was tested, showing a concentration-dependent increase in blebbing and actin cortex formation, with 75 µM identified as the optimal concentration. Control cells were treated with the vehicle (DMSO) at the same final concentration as in the corresponding ionomycin-treated samples.

### Ruthenium red

0.25 M ruthenium red (Abcam, ab120264) was added to cell cultures. Samples were incubated for 5 min before imaging. Control cells were treated with the vehicle (H2O) at the same final concentration as the ruthenium red-treated samples.

### Microvilli ablation

To ablate *S. rosetta* microvilli, glycocalyx digestion was performed using papain [43]. This treatment removes microvilli while preserving overall cell integrity. A papain working solution was prepared by diluting 5 μL of commercial papain (Sigma-Aldrich, P3125-100MG) in 45 μL of papain dilution buffer (50 mM HEPES-KOH, pH 7.5, 200 mM NaCl, 20% (v/v) glycerol, and 10 mM L-cysteine; 0.22 μm filtered and stored at −80°C). Subsequently, 4 μL of the diluted papain solution were mixed with 400 μL of digestion buffer (40 mM HEPES-KOH, pH 7.5, 34 mM lithium citrate, 50 mM L-cysteine, and 15% PEG 8000; 0.22 μm filtered and stored at −80°C), resulting in a final papain concentration of 1 μM.

Cells were incubated in the papain solution for 35 min at room temperature. Digestion was stopped by adding 10 μL of 50 mg/mL bovine serum albumin (BSA; Fisher Scientific, 12877172), prepared in distilled water. Artificial seawater (ASW) was then added to a final volume of 1 mL. Cells were subsequently confined and imaged within 15 min of terminating the digestion, before microvilli regeneration occurred.

### Genetic modification: production of the kikGR-actin endogenously tagged strain

Codon-optimized kikGR and mEOS sequences for *S. rosetta* were designed using Geneious Prime 2026.1.2, based on published protein sequences [37,84] and on *S. rosetta* codon usage for highly expressed genes [43]. Codon-optimized sequences were ordered as gene fragments from Integrated DNA Technologies (IDT) and cloned using restriction enzymes (ClaI and BsiwI) into a previously published expression vector for *S. rosetta* under an actin promoter and with a 3’UTR from *S. rosetta* EF1α, that additionally contains a Pac puromycin resistance gene [11]. The resulting plasmid (pTB25) was transfected and puromycin selection was performed as in [48]. Examination of cells under a Leica Stellaris 8 confocal microscope did not reveal significant fluoresence in mEOS-expressing cells, but green cytoplasmic fluorescence was noticed in kikGR-expressing cells, and could be photoconverted into red fluorescence following 10 rounds of exposure to a 405 nm laser at 10% power. This was taken as an indication that kikGR can serve as a photoconvertible marker in *S. rosetta*.

We then produced a plasmid encoding a version of *S. rosetta* actin tagged at its N-terminal end with kikGR followed by the flexible linker GSSGSSGSSGSS (encoded by the nucleotide sequence 5’-GGCTCCTCTGGTAGCTCCGGCTCTAGCGGTTCTTCC-3’). *S. rosetta* has three actin loci encoding identical predicted protein sequences (PTSG_05705, PTSG_01553, PTSG_02201). PTSG_01553 was amplified from *S. rosetta* RNA (extracted as in [40]) by one-step RT-PCR using the SuperScript IV One-Step RT-PCR kit (ThermoFisher 12595025) with primers CC86 (CTTCCATGGGTGACGAGGATGTTGC) and CC87 (CTCATCGATTTAGAAGCACTTGCGGTGGACAA). RT was performed for 10’ at 50°C and reverse transcriptase was inactivated for 2’ at 98°C. PCR was performed using the following program: 5x (98°C 10”, 58°C 10”, 72°C 45”); 30x (98°C 10”, 70°C 10”, 72°C 45”); 5’ at 72°C. The PCR product was validated by Sanger sequencing (Eurofins Genomics) and amplified further using DreamTaq (ThermoFisher Scientific EP0711).

The linker sequence was then added to pTB25 by PCR using primers CC85 (CACATCGATCCATGGAAGAACCGCTAGAGCCGGAGCTACCAGAGGAGCCGGCCTCGAACTCGTACTTGG) and CC65 (CACAAAACAACCAGCCGTACGATGTCCGTCAT). In parallel, the actin-encoded PCR product was digested using NcoI and ClaI, and inserted by restriction cloning in the pTB25+linker amplicon.

The resulting plasmid was used as a repair template for recombination with the endogenous actin locus PTSG_05705, using a previously reported gene replacement strategy [40]. The two following guide RNAs were used to remove the endogenous gene: PTSG_05705_gRNA1 (ACCACAAAACAACCAGCCAT) and PTSG_05705_gRNA2 (CAAGGTGGCAATGACCAACG). The repair template carried 50 bp homology arms, kikGR-actin, and a puromycin resistance cassette. Transfection, puromycin selection and clonal isolation were performed as in [40]. Clones were genotyped by PCR using Q5 high-fidelity DNA polymerase (NEB M0491L) using the primers CC103 (CTGAAAAGCCAATGCCGTCC) and CC104 (TGTTGTCGCAGCTCTACGTT). Cells expressing the kikGR-actin fusion construct were imaged and photoconverted with the same protocol as those expressing kikGR alone (see above).

### Genetic modification: endogenous C-terminal fluorescent tagging of ERM1

To visualize ERM1 protein in live cells, we used a protocol adapted from [40,41] to generate an ERM1::StayGold strain (Pillai et al, in preparation). This resulted in an in-frame insertion of the mStayGold fluorescent marker at the 3’ end of the gene coding sequence to generate an endogenously tagged C-terminal fusion protein in *S. rosetta*.

We designed a crRNA (protospacer 5’-AGAGCATGTAACCAGCAACA-3’; PAM 5’-CGG-3’) targeting the *erm1* locus in the 3’UTR, 6 bp downstream of the stop codon. The repair template was amplified by PCR from the pMD01 plasmid, which carries a codon-optimized mStayGold coding sequence preceded by a flexible linker (GSGSGGSGSGSGGSGS, nucleotide sequence 5’-GGGTCGGGAAGTGGCGGCAGGGAAGTGGGAGTGGAGGGAGCGGTTCT-3’) as well as a puromycin-resistance cassette. Sequence for the linker and for codon-optimized mStayGold were previously reported [48].

Repair templates (linker-mStayGold-EF1 3’UTR followed by puromycin-resistance cassette) were produced by PCR with ∼40-50 basepair homology arms framing the CRISPR/Cas9 cut site in the *erm1* locus:

- Forward primer EP83: 5’-CATTCGCGCCGGAAACACACGCCAACGCATCCTTGACTTTGAGAGCATGGGGTCGGGAAGTGGCGGCAGCGGAA-3’
- Reverse primer EP84: 5’-CAAAACAGACATACACATTCACAGCCACGCATGCGTATAGGTGTTGCTGGAAGCTTGTAACGACTGCAGTCTTGT-3’

The forward primer was designed with a 49 bp 5′ homology arm corresponding to the final 49 bp of the *erm1* coding sequence immediately before the native stop codon. This design removed the endogenous stop codon and generated an in-frame fusion of the coding sequence with the linker sequence encoded by the repair template. The reverse primer contained a 38 bp homology arm. During homology-directed repair, recombination between the forward and reverse homology arms resulted in deletion of the intervening 15 bp of native sequence, encompassing the native stop codon.

Repair template amplification, purification, co-transfection with the Cas9:gRNA ribonucloprotein, selection and clonal isolation were performed as in [40]. Prior to confocal imaging, the plasma membrane of cells expressing ERM1-mStayGold was stained by adding BioTracker MemBright 640 (Sigma Aldrich, SCT085) at 1:10,000 dilution.

### Genetic modification: production of the KRas-mStayGold-expressing strain

The KRas-mStayGold strain was generated using a crRNA (CATGGACGAGGAAGCAAGGA) specific to the *ura3* gene (PTSG_03344; generous courtesy of David S. Booth, University of California, San Francisco). Custom primers containing a termination cassette and 50 bp homology arms, MA124 (CACCACCAACCAACAAACCAACCAACCACCAACCATGGACGAGGAAGCAATTTATTTAATTAAATAAACAAACTGCCCTGCACCTCGTCTGC) and MA125 (GCCAAACTTGACGGCCCCAATGTCAAACAGCCGAAGCACAAGCGCCTTCCTTTATTTAATTAAATAAAACAGATACCAACGTGTCGAGTC) were purchased from Merck Sigma-Aldrich and used to amplify a published plasmid template encoding KRas-mStayGold tag [48]. Two PCR reactions were set up in 50 μL using Q5 high fidelity DNA polymerase (New England Biolabs M0491S) with 100 ng plasmid as template, 200 μM dNTP (ThermoScientific R1121), 0.5 μM of each primer, and 0.02 U/μL Q5 polymerase. The following PCR program was run on a Biorad PTC Tempo 96 Thermal Cycler: 30’’ 98°C; 40X (10’’ 98°C; 45’’ 67°C; 2’ 72°C); 2’ 72°C; hold 10°C. Size of a PCR products (expected: 3,6 kb) was visually checked by running 1:10 of the PCR reaction product on a Blue-Light Transilluminator (Invitrogen G6600EU). The two PCR products were combined and purified using a PCR purification kit (Macherey NucleoSpin Gel and PCR cleanup 740609.50) and the final product was eluted with 17 μL Elution buffer. 2 μL were used to measure DNA concentration using a NanoDrop spectrophotometer (LabTech). The remaining 15 μL were reconcentrated in 2 μL by evaporation for 1 hour at 55°C and then used as repair template for nucleofection (4.2 μg). Cell recovery post nucleofection and puromycin selection were performed at 30°C instead of 25°C. After clonal selection, a genotyping PCR reaction was set up with Q5 Polymerase following provider specifications, with 1 μL genomic DNA as template, 200 μM dNTP, 0.5 μM each primer (MA128_FW: TCTGCCTTGTGCCATGGGCTG and MA128_REV: GACGCCACACAGGGTGTCATA) and 0.02 U/μL Q5 polymerase. The following PCR program was run: 30’’ 98°C; 40X (20’’ 98°C; 30’’ 71°C; 2’ 72°C); 4’ 72°C; hold 10°C. PCR product was run on gel, purified as above, and sequenced by Plasmidsaurus (Oxford Nanopore, R10.4.1).

### Cell Segmentation and morphometrics for image analysis of Ca^2+^ signaling data

Cell contours were segmented from the DIC channel using CellPose [79]. A custom model was generated by retraining the cyto3 model with representative images from the dataset to improve segmentation accuracy. The trained model was subsequently employed to process time-lapse recordings through TrackMate [85], enabling automated tracking of individual cells over time.

Following segmentation, binary masks were generated for each frame and used to extract morphometric parameters. To quantify blebbing activity, temporal changes in cell shape were analyzed by computing the frame-to-frame variation in projected cell area for each tracked cell, as in [11]. The blebbing activity index was defined as the absolute difference in cell area between consecutive time points. Custom scripts are available at [86].

### Ca^2+^ pulses quantification

Time-lapse fluorescence recordings were acquired using the RGECO channel under identical imaging conditions for all samples. Single Z images were taken with a 63x water immersion C-Apochromat 1.1 NA objective mounted on a Zeiss Observer Z.1. For cell segmentation, summed intensity projections over the whole timelapse were first generated in Fiji [87,88] using the RGECO channel to improve signal-to-noise ratio. Cells were segmented using Cellpose version 2.3 [79] with the cyto3 model as a base, followed by manual correction to ensure accurate boundaries.

For visualization and extraction of fluorescence traces, RGECO intensity values were measured over time for each segmented cell using Fiji’s Measure function. The resulting time-series data were exported as CSV files and further processed in Python (version 3.10) using custom scripts.

Fluorescence traces were normalized to their baseline intensity (F_0_, defined as the 10th percentile of each trace), and ΔF/ F_0_ was calculated for each time point

Automatic peak detection was performed using the find_peaks function in SciPy (minimum prominence = 2 SD). Peak-derived features—including amplitude, width, area, and inter-peak intervals—were extracted for each cell. Custom scripts are available at [89].

### Tomography

#### Sample preparation for cryo-ET

Before freezing, *S. rosetta* cultures were scaled up to 2 L, centrifuged at 2000 × g for 15 min at room temperature, and washed three times with ASW (2500 × g, 5 min each) to remove excess *E. pacifica*.

#### Plunge freezing/vitrification of amoeboid *S. rosetta*

*S. rosetta* cells were pelleted and resuspended in a small volume of artificial seawater (ASW; 20 µL) immediately prior to freezing. The concentrated cell suspension was applied onto cryo-EM copper or gold grids with a holey silicon film (Quantifoil SiO₂ R2/2, 200 mesh). Prior to sample application, grids were placed on a glass slide and plasma-cleaned for 1 min at 15 mA and 0.38 mbar using a PECO easiGlow system. 1 µL of samples was then transferred to a Leica EM GP2 automated plunge freezer (adding 2 µL of ASW at the back of the grid), equilibrated to 20 °C and 98% relative humidity, and back-blotted for 9 s using Whatman Grade 1 filter paper to remove excess buffer. Grids were immediately plunge-frozen into liquid ethane (−180 °C).

Vitrified grids were mounted into Autogrids (Nanosoft) with notched rings for FIB milling, secured using C-clips (Thermo Fisher Scientific, Cat. No. 1036171) and stored in liquid nitrogen until further use.

#### High Pressure Freezing

Cells were high-pressure frozen using a Leica ICE system (Leica Microsystems). Small aliquots of concentrated *S. rosetta* cultures were stained with FM 4-64 (Thermo Fisher, T13320) at a 1:10,000 dilution, followed by addition of *E. pacifica* food pellet diluted 1:100.

For waffle preparation, cryo-EM gold grids with a holey silicon film (Quantifoil SiO₂ R2/2, 200 mesh) were embedded in the sample. Grids were placed into 6 mm type B carriers (Wohlwend GmbH) coated with soy lecithin (15 mg/mL in chloroform; Sigma-Aldrich 319996) and glow-discharged for 1 min at 25 mA.

For lift-out preparation, 4 µL of sample was placed onto 3 mm type A (50 µm or 100 µm) carriers (Wohlwend GmbH) and covered with 3 mm type B carriers, similarly coated with soy lecithin (15 mg/mL in chloroform; Sigma-Aldrich 319996) and glow-discharged for 1 min at 25 mA. Planchettes were assembled and subjected to high-pressure freezing at ∼2000 bar, vitrifying the samples. Frozen specimens were transferred under liquid nitrogen and stored in liquid nitrogen until further processing.

#### Cryo-confocal imaging

Samples were imaged using either a cryo-widefield Leica THUNDER Imager EM Cryo CLEM, with a Leica HCX PL APO 50x / 0.90 CLEM (Leica Microsystems) objective or a cryo-Stellaris 8 confocal system (Leica Microsystems) equipped with a HC PL APO 50×/0.90 C cryo objective, a white light laser (WLL), and three hybrid (HyD) detectors. A two-dimensional widefield overview was initially acquired to evaluate grid integrity, ice quality, and the spatial distribution of cells in regions suitable for imaging. Subsequently, cryo-confocal image stacks were collected with the pinhole set to 1 Airy unit (AU), a zoom factor of 1.7×, and a z-step size of 0.5 µm.

#### Cryo FIB miling

Autogrids containing vitrified *S. rosetta* cells were cryogenically transferred into an Aquilos 2 dual-beam FIB/SEM instrument (Thermo Fisher Scientific) equipped with a cryo-stage precooled to −185 °C. To improve electrical conductivity, a thin platinum layer was first applied using magnetron sputter coating (1 kV, 30 mA, 60 s), followed by an additional platinum deposition using the gas injection system (GIS) preheated at 28°C for XXs to protect the sample surface from irregular milling side effect.

A grid overview was acquired in SEM mode, and suitable cells were identified using MAPS v3.29 software (Thermo Fisher Scientific). For milling, the stage was tilted to a 10° milling angle between the grid plane and the gallium ion beam. Cryo-FIB milling was performed using a gallium ion beam with progressively decreasing currents—typically 0.3 nA for coarse milling, 0.1 nA for thinning, and 30 pA for final polishing. Stress-relief cuts (0.6 µm × 8 µm × 7 µm) were introduced 4 µm from the lamellae to minimize mechanical strain. This procedure yielded self-supporting lamellae approximately 150 nm thick. SEM imaging was conducted at 2–5 kV and 13–20 pA to monitor milling progression.

For samples prepared by high-pressure freezing, either the Waffle Method or Serial Lift-Out was used for lamella preparation as previously described in [51] and [50], respectively. In both cases, integrated fluorescence light microscopy (iFLM) was directly used on Aquilos to identify regions of interest prior to cryo-FIB milling.

#### Titan Krios Acquisition

Cryo-electron tomography was performed on cryo-FIB–milled lamellae using a Titan Krios transmission electron microscope (Thermo Fisher Scientific) operated at 300 kV and equipped with a cold field emission gun (C-FEG), a Falcon 4i direct electron detection camera (Thermo Fisher Scientific), and a Selectris X energy filter (Thermo Fisher Scientific) operated in counting mode. Microscope control and data acquisition were performed using Tomography 5 software v5.6 (Thermo Fisher Scientific).

The microscope was operated in nanoprobe mode with the energy filter slit width set to 10 eV, the C2 aperture to 70 µm, and the objective aperture to 100 µm. Dose-symmetric tilt series (Hagen et al., 2017) were collected from +70° to −50°, starting with a 10° offset to compensate for lamella pre-tilt, with 2° increments. Data were acquired at a nominal magnification of 42,000×, corresponding to a pixel size of 3.101 Å. The defocus range was set between −3 and −5 µm, and the total accumulated dose was limited to 140 e⁻/Å².

Image frames were recorded in EER format. The cold field emission gun was flashed prior to each acquisition to adjust dose rate and then software-controlled.

In total, 40 tilt series were acquired from *S. rosetta* cells in the amoeboid state, 40 in the transition state, and 20 in the flagellate state.

#### Tilt series alignment and reconstruction

Tilt movie frames were motion-corrected using the Scipion framework [90]. Tilt series alignment was performed with AreTomo version 2 [91] within the Scipion interface. The aligned tilt series were reconstructed in AreTomo using weighted back-projection combined with a SIRT-like filter (10 iterations). Reconstructed tomograms were subsequently binned to a final pixel size of 12.4 Å and denoised using Cryo-CARE [92]. Some of the lift-out tomograms were manually reconstructed using IMOD and denoised using CryoSAMBA [93]. We noticed at this step that ice quality appeared lower in high-pressure frozen samples than in plunge-frozen samples, presumably due to less efficient vitrification of thick samples despite high-pressure freezing.

#### Tomogram segmentation and template matching

Selected tomograms were used as input for the automated MemBrain segmentation pipeline [94] employing the v10_alpha pre-trained model. Microtubule segmentation was subsequently performed using Easymode [95]. The resulting segmentations were manually curated in Napari [96] to correct local misassignments and ensure structural continuity of cellular and subcellular features. Manual validation was guided by visual inspection of the corresponding tomographic slices taken from 3D reconstructions. For visualization and movie generation, the curated segmentations were imported into UCSF ChimeraX v1.9 [97].

#### Actin Segmentation and quantification

Actin filaments were manually segmented in 3D from denoised cryo-electron tomograms (voxel size 12.4 Å) using the brush tool in Napari, producing binary volumetric masks.

Cortical actin density was quantified per membrane mesh vertex using a custom napari plugin developed for this study and available for installation via pip [98]. Input meshes were triangulated surface meshes of the plasma membrane generated with the surface morphometrics pipeline ([99], and actin was provided as a binary volumetric segmentation in the same tomogram coordinate space, which the plugin skeletonizes internally prior to density calculation.

For each mesh vertex, we counted skeletonized actin filament points falling within a cylindrical search volume oriented along the inward-pointing surface normal, with a height of 200 nm and a lateral radius of 100 nm. Actin filament counts were converted to local density in units of µm of filament per µm² of membrane surface:

- density (µm/µm²) = (actin skeleton points × voxel size [nm] × 10⁻³) / (π × lateral radius² [nm²] × 10⁻⁶)

Surface normals, whose initial orientation from mesh triangulation is arbitrary, were oriented inward using nearby actin skeleton points as a local reference where sufficient actin was present nearby, with orientation propagated to the remaining vertices via breadth-first search along mesh connectivity, computed independently within each connected mesh component to prevent cross-contamination between membranes in multi-component meshes.

Density values were visualized directly on the 3D membrane mesh in napari as a turbo colormap heatmap, with contrast limits set to the 5th–95th percentile of the density distribution to reduce sensitivity to outlier vertices. To provide quantitative reference for the heatmap, a matched colorbar indicating the density scale was generated, and the underlying per-vertex density distribution was plotted as a histogram to assess the shape and range of local variation across the membrane surface. Per-vertex density values, mesh coordinates, and pipeline parameters were exported as .csv and .json files alongside a density-mapped mesh for downstream statistical comparison and figure rendering. The script used for actin density quantification was deposited at [98].

#### Actin skeleton segmentation and preprocessing

Actin filaments were manually segmented in 3D from denoised cryo-electron tomograms (voxel size 12.4 Å) using the brush tool in napari, producing binary volumetric masks. Masks were skeletonized in 3D (skimage.morphology.skeletonize), and skeletons were broken at branch points using a custom 18-connectivity kernel restricted to within-slice contacts (excluding cross-Z-slice diagonal connections), which was empirically validated against direct tomogram inspection to avoid spurious fusion of adjacent, non-continuous filaments. Connected components smaller than 4 voxels after branch-breaking were discarded as noise. Each remaining connected component was treated as an individual filament fragment for downstream analysis.

#### Bundle vs. mesh classification

Filament fragments were classified as belonging to a bundle (tightly packed, parallel actin) or to the surrounding mesh (isolated/crossing filaments), following the operational criterion of [100]: two actin points are considered a bundle contact if they lie within a threshold distance and their local orientations differ by less than a threshold angle. Local filament orientation at each skeleton point was estimated by local PCA over a small spatial window restricted to points of the same fragment. Candidate neighbor pairs across different fragments were identified using a KD-tree (scipy.spatial.cKDTree.query_pairs), with an additional perpendicularity filter requiring that the displacement between paired points be predominantly lateral (not along the filament’s own axis), to avoid mistaking longitudinal continuations of a single filament for a genuine inter-filament contact. For each point, a bundle contact was scored if it had a neighbor from a different fragment satisfying both the distance and angle thresholds. Each filament fragment was then classified as bundle if ≥50% of its points scored as bundle contacts (majority vote).

Distance and angle thresholds were selected via a sensitivity sweep over distance (10, 15, 20 nm), angle (15°, 20°, 25°), and a minimum-parallel-neighbor count N (1–5), for 45 total parameter combinations. The most stringent combination tested (10 nm / 15° / N ≥ 5) was selected as final, based on visual agreement in napari between classified bundle regions and regions of visually dense, parallel filament packing. The script used for this analysis is available at [101]

#### Inter-filament spacing within bundles

For filament pairs within classified bundles, center-to-center spacing was measured as follows. Filament coordinates (in nm) were divided into Z-slices; within each slice, a line was fit to the XY coordinates of each filament by singular value decomposition (SVD). Neighboring filament pairs were fixed globally based on 3D centroid proximity (with a filter excluding pairs that had another filament positioned between them), using a maximum neighbor distance of 35 nm. For each Z-slice in which both filaments of a pair were present, center-to-center distance was measured as the perpendicular distance between their fitted lines in the XY plane. Surface-to-surface gap was calculated as center-to-center distance minus twice the filament radius (3.5 nm, corresponding to the ∼7 nm diameter of F-actin). The script used for this analysis is available at [102].

#### Visualization

For publication figures, density-mapped meshes, bundle/mesh classification volumes, and the actin skeleton were exported in BILD format (the format found to reliably preserve per-face/per-vertex color when imported into ChimeraX, as VTP and PLY export/import did not preserve color mapping in our ChimeraX version) and rendered in ChimeraX. Density heatmaps used a turbo colormap; bundle/mesh classification used a colorblind-safe palette (Okabe-Ito), with binary volumes exported per classification category.

#### Figures and statistics

All data analysis and figure generation were performed using Python (v3.10), and statistical analyses were carried out within the same environment. Final figure layouts and graphical assembly were completed in Adobe Illustrator v29.1 (Adobe Inc.).

## Acknowledgements

We thank Debbie Maizels for assistance with the figures; Jeff Colgren and Pawel Burkhardt for sharing the RGECO-expressing strain of *S. rosetta*; David S. Booth for the sequence of the ura3 guide RNA; Maxwell Coyle, Alain Garcia de las Bayonas and Nicole King for feedback on the manuscript; Mohamad Harastani for help with Scipion; Mart So-Last for help with EasyMode; Erwan Fournery for introducing the PHEM fixation buffer; Alba Diz-Muñoz, Detlev Arendt, Lucas Leclère, Pablo Vargas, Matthieu Piel, Ana-Maria Lennon-Duménil, and all members of the ECB lab for useful suggestions and feedback during the project. Panels illustrating cryofixation in Figure 4 were created using BioRender.com. M.F.-D. is supported by a doctoral fellowship from Université Paris cité (École doctorale Bio Sorbonne Paris Cité, BioSPC, ED 562) and a 4^th^ year fellowship from the Fondation pour la Recherche Médicale. M. Ansel is supported by the Ecole Normale Supérieure de Lyon (allocation élève normalien). E.P. was supported by an EIPOD4 fellowship (this project has received funding from the European Union’s Horizon 2020 research and innovation programme under the Marie Skłodowska-Curie grant agreement No 847543), an Add-on Fellowship for Interdisciplinary Science from the Joachim Herz Stiftung, and a Company of Biologists Travelling Fellowship. We acknowledge the cryo-ET expertise and assistance of the Institut Pasteur’s NanoImaging Core facility, created and supported by a PIA grant (EquipEx CACSICE: ANR-11-EQPX-0008). UTechS PBI is part of the France–BioImaging infrastructure network (FBI) supported by the French National Research Agency (ANR-10-INBS-04; Investments for the Future), and acknowledges support from Institut Pasteur, ANR/FBI, the Région Ile-de-France (DIM1HEALTH) and the French Government Investissement d’Avenir Programme— Laboratoire d’Excellence “Integrative Biology of Emerging Infectious Diseases” (ANR-10-LABX-62-IBEID). T.B. and his laboratory are supported by the Institut Pasteur (G5 package), the European Research Council (EvoMorphoCell, grant no. 101040745), the Vallee Foundation, the EMBO Young Investigator Programme, the LabEx Revive (ANR-10-LABX-73), the Agence Nationale de la Recherche (INCOMPLETE, ANR-23-CE13-0031) and the CNRS (UMR 3691). Views and opinions expressed are those of the author(s) only and do not necessarily reflect those of the European Union or the European Research Council. Neither the European Union nor the granting authority can be held responsible for them.

## Author contributions

Maite Freire-Delgado designed the project, acquired funding, performed most experiments, prepared figures and edited the manuscript. Diede de Haan contributed to cryo-EM experiments alongside M. F.-D. Eva K. Pillai designed and produced the ERM1-mStayGold strain and performed the experiment involving it alongside M. F.-D. Mylan Ansel performed UExM, generated some of the corresponding figure panels, and generated the KRas-mStayGold strain used in the study. Stéphane Tachon and Anastasia Gazi contributed to cryo-EM experiments alongside M. F.-D. Marvin Albert designed the image analysis pipeline for cortex and blebbing quantifications, performed the corresponding analyses, and produced graphs. Chantal Combredet generated the kikGR-actin strain. Audrey Salles performed SIM alongside M. F.-D. and T. B. Jean-Yves Tinevez supervised M. Albert. Thibaut Brunet designed and supervised the project, acquired funding, acquired preliminary data, prepared figures and wrote the manuscript. All authors read, edited and approved the final manuscript.

---

**Figure S1.**
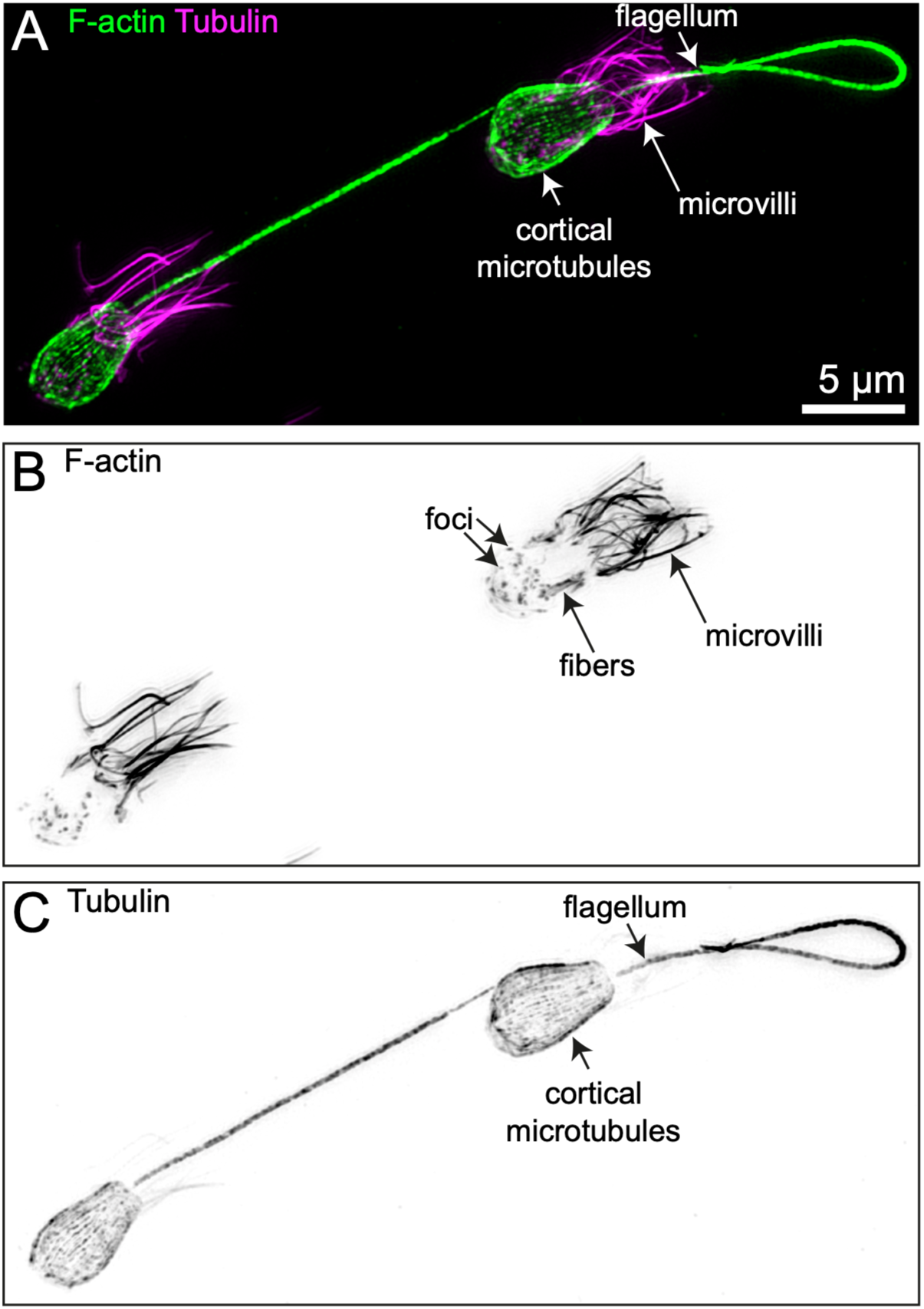
Structured Illumination Microscopy reveals the cytoskeletal ultrastructure of the flagellate form of *S. rosetta*. Immunostaining confirms the presence of microvillar F-actin, F-actin foci in the cell body (plausibly endocytic patches), cortical microtubules, and flagellar tubulin. No F-actin cortex is detectable.

**Figure S2.**
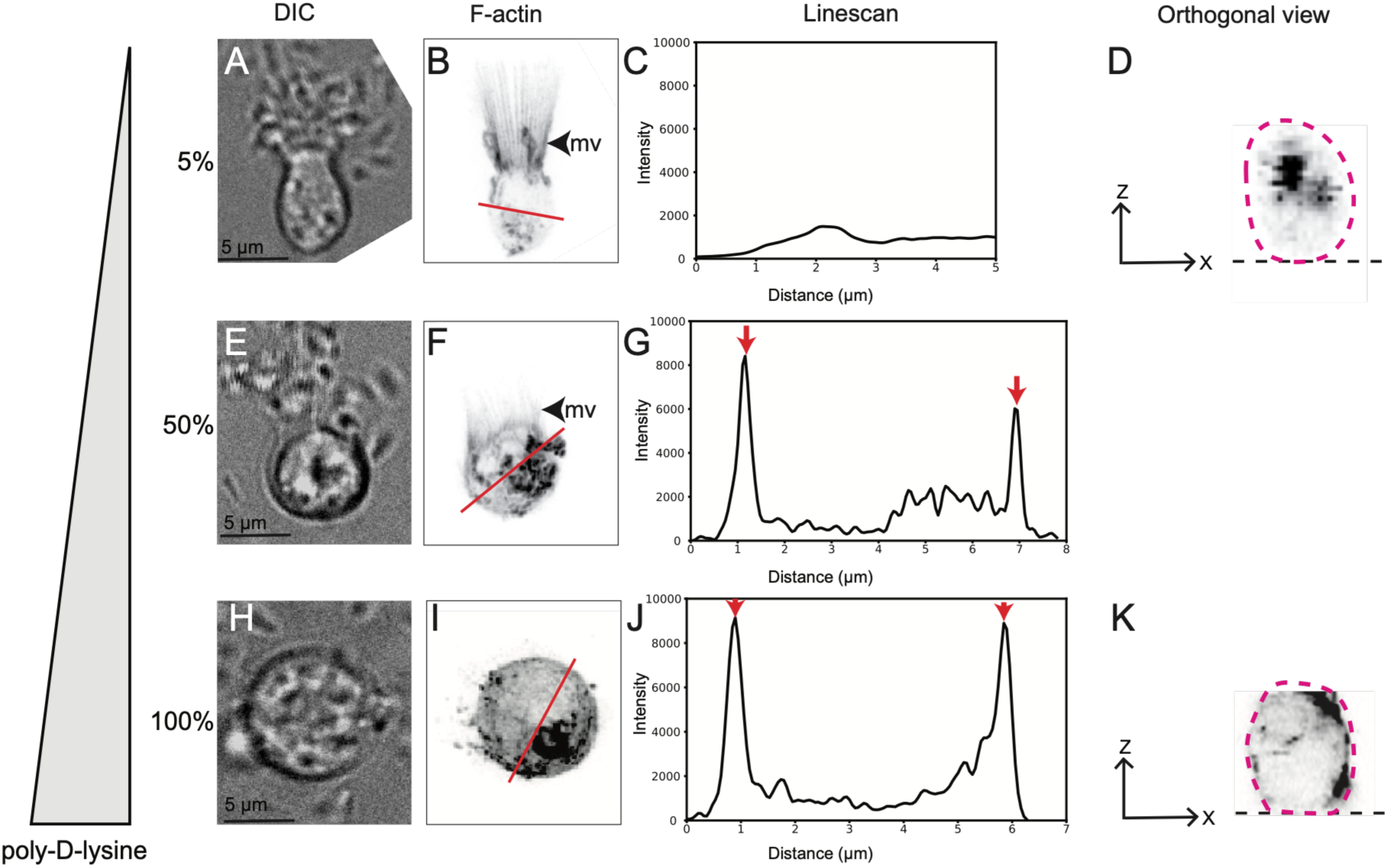
Poly-D-lysine induces cortex formation at high concentration. Cells were plated on wells coated with different concentrations of poly-D-lysine, prepared by dilution of a 10 mg/mL stock in distilled water. After coating, liquid was removed and cells were plated on the coated well without any additional wash. (A-D) Low concentrations of poly-D-lysine (5%) resulted in cell immobilization, but preserved the native flagellate architecture, as characterized by a regular microvillar collar and no actin cortex. (A) Transmitted light micrograph. (B) F-actin signal (LifeAct-mCherry). (C) Linescan along the line in B. (D) Virtual cross-section along the y axis, showing that the cell body (red dotted outline) retains an ovoid shape and is not flattened at the contact of the well (black dotted line). (E-G) Intermediate concentrations (50%) induce a partial amoeboid transition: microvilli remain visible but are shortened and a cortex appears. (E) Transmitted light, (F) LifeAct-mCherry fluorescence, (G) Linescan across the red line in F. Red arrows: cortex. (H-K) High concentrations (H-K) induce a complete amoeboid switch, with retracted collar and strong cortex. (H) Transmitted light, (I) LifeAct-mCherry, (J) Linescan across the red line in I (red arrows: cortex), (K) Virtual section across the y-axis, showing flattening of the cell body (dotted red outline) where it contacts the bottom of the well (black dotted line).

**Figure S3.**
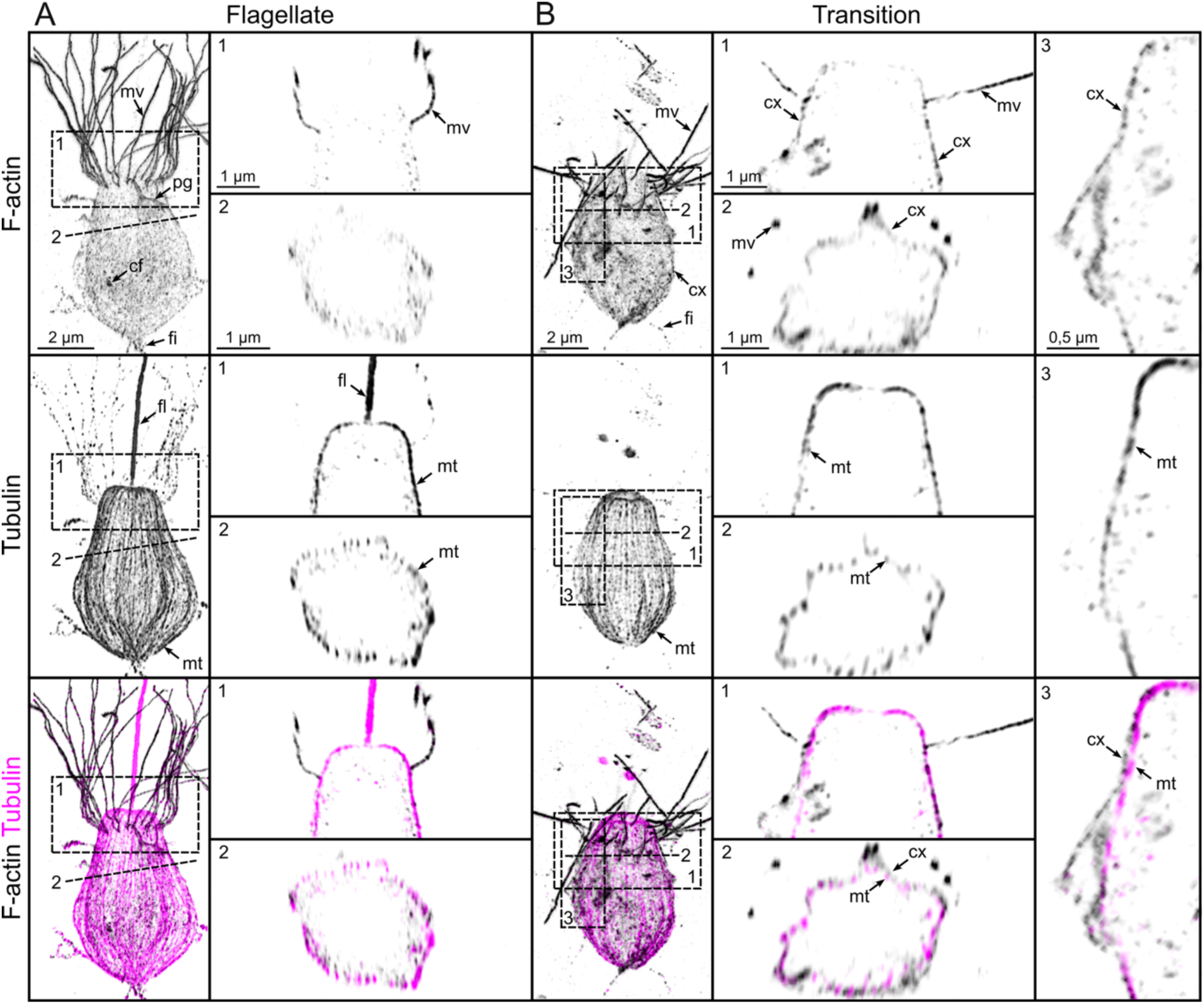
Ultrastructural expansion microscopy (U-ExM) highlights F-actin cortex formation in the transition state. (A) Representative flagellate cell. The cell displays a tubulin-labeled apical flagellum, cortical microtubules, F-actin-rich apical microvilli, cytoplasmic actin foci (putative endocytic patches), a phagocytic cup, and basal filopodia. Notably, no actin cortex is visible (see longitudinal and transversal cross-sections 1 and 2). (B) Representative transition-state cell lacking its flagellum. The cell shows partly retracted microvilli and an incipient F-actin cortex, while cortical microtubules remain present. Continuous actin filaments extend from the microvilli to the nascent cortex, suggesting recruitment of the microvillar actin pool (longitudinal cross-section 1). The incipient actin cortex forms above the cortical microtubules, potentially replacing them as the primary cortical cytoskeleton (transversal and longitudinal cross-sections 2 and 3). Cells were labeled with β-tubulin (magenta) and HAK-actin (black). In all panels: cf, cytoplasmic foci; cx, cortex; fg, flagellum; fi, filopodia; mv, microvilli; mt, microtubules; pg, phagocytic cup.

**Figure S4.**
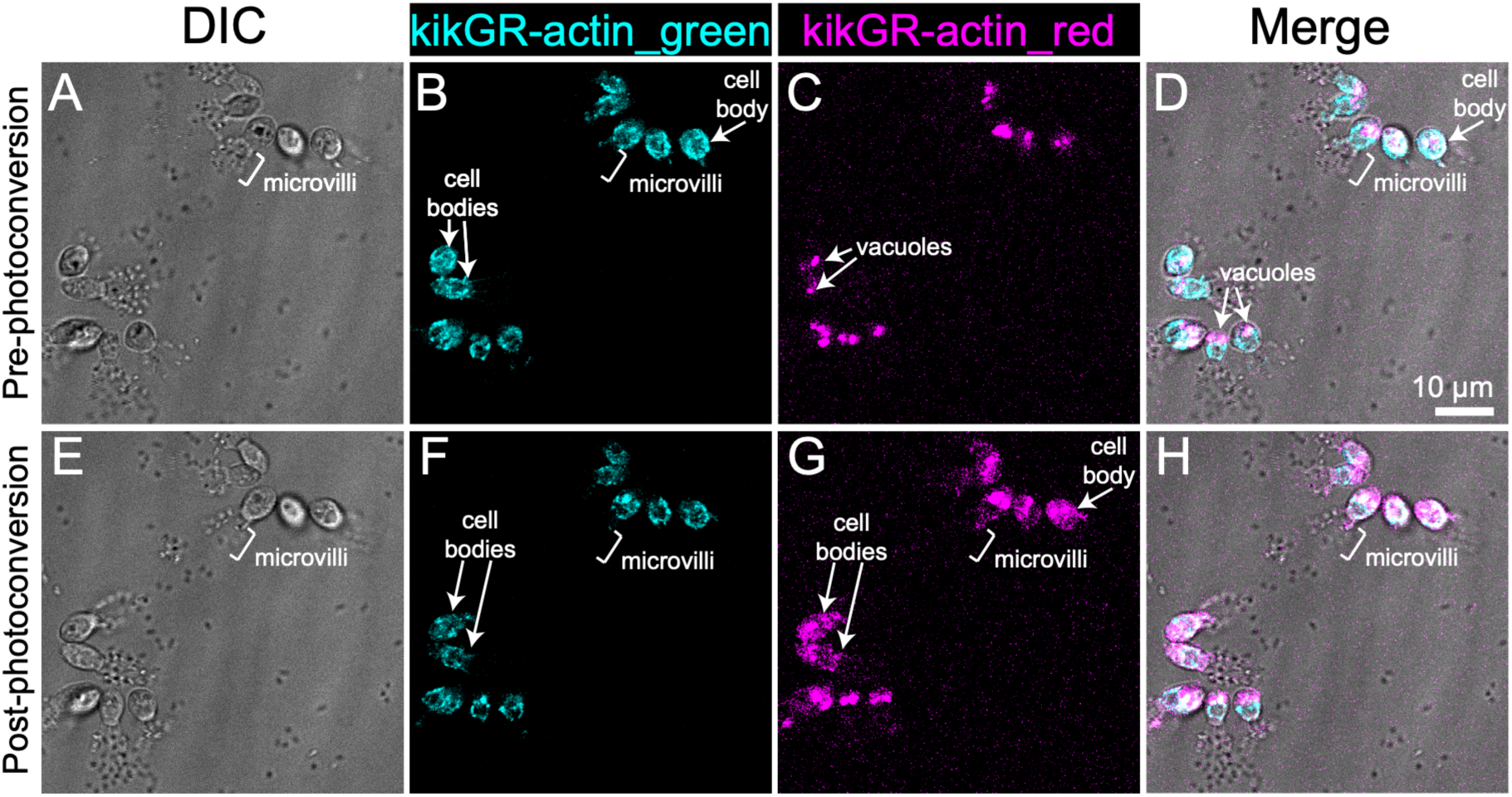
kikGR-actin is photoconvertible but does not incorporate in microvilli. (A-D) Before photoconversion, green kikGR-actin is detected in the cell body, but not in microvilli. Red fluorescence is restricted to autofluorescent vacuoles, as previously reported [103]. (E-H) Photoconversion converts part of the green cytoplasmic fluorescent signal into red cytoplasmic fluorescent signal, but microvilli remain non-fluorescent.

**Figure S5.**
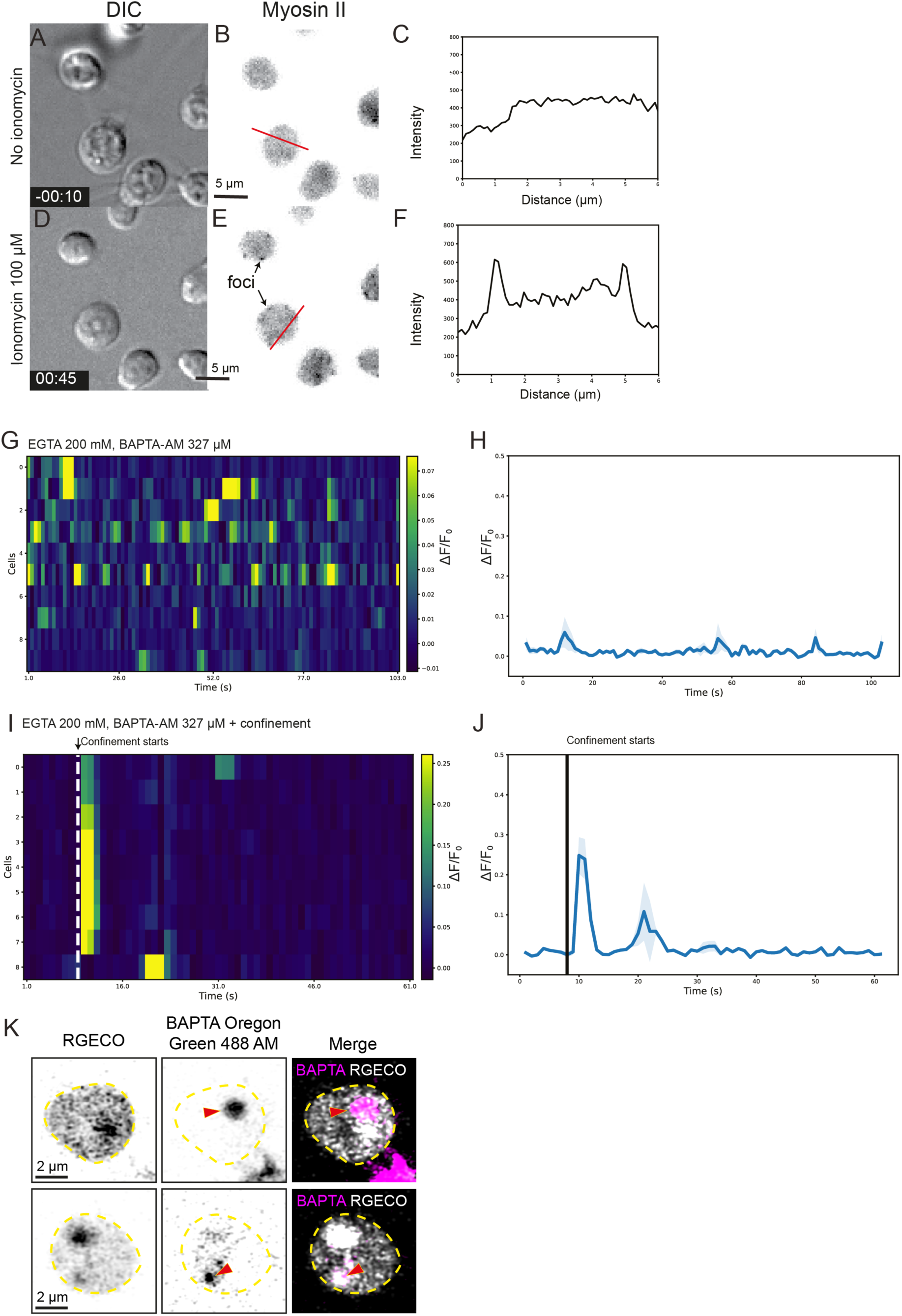
Calcium signaling in the amoeboid switch. (A-C) In negative control cells, myosin II is diffusely present in the cytoplasm. (A) Transmitted light channel. (B) MRLC-mTFP. (C) Linescan along the red line in B. (D-F) Upon ionomycin treatment, myosin II condenses into cortical foci and fibers, as previously described upon confinement [11]. (D) Transmitted light. (E) MRLC-mTFP. (F) Linescan along the red line in E. (G-H) Treatment with extracellular (EGTA) and intracellular (BAPTA-AM) calcium chelators does not abolish spontaneous calcium pulses, as visualized by RGECO fluorescence. (I-J) Treatment with extracellular (EGTA) and intracellular (BAPTA-AM) calcium chelators does not abolish confinement-induced calcium signaling, as visualized by RGECO fluorescence. (K) The BAPTA-AM fluorescent analog BAPTA Oregon Green 488 AM accumulates in vacuoles but is indetectable in the cytoplasm, indicating that *S. rosetta* can sequester BAPTA derivatives in intracellular compartments.

**Figure S6.**
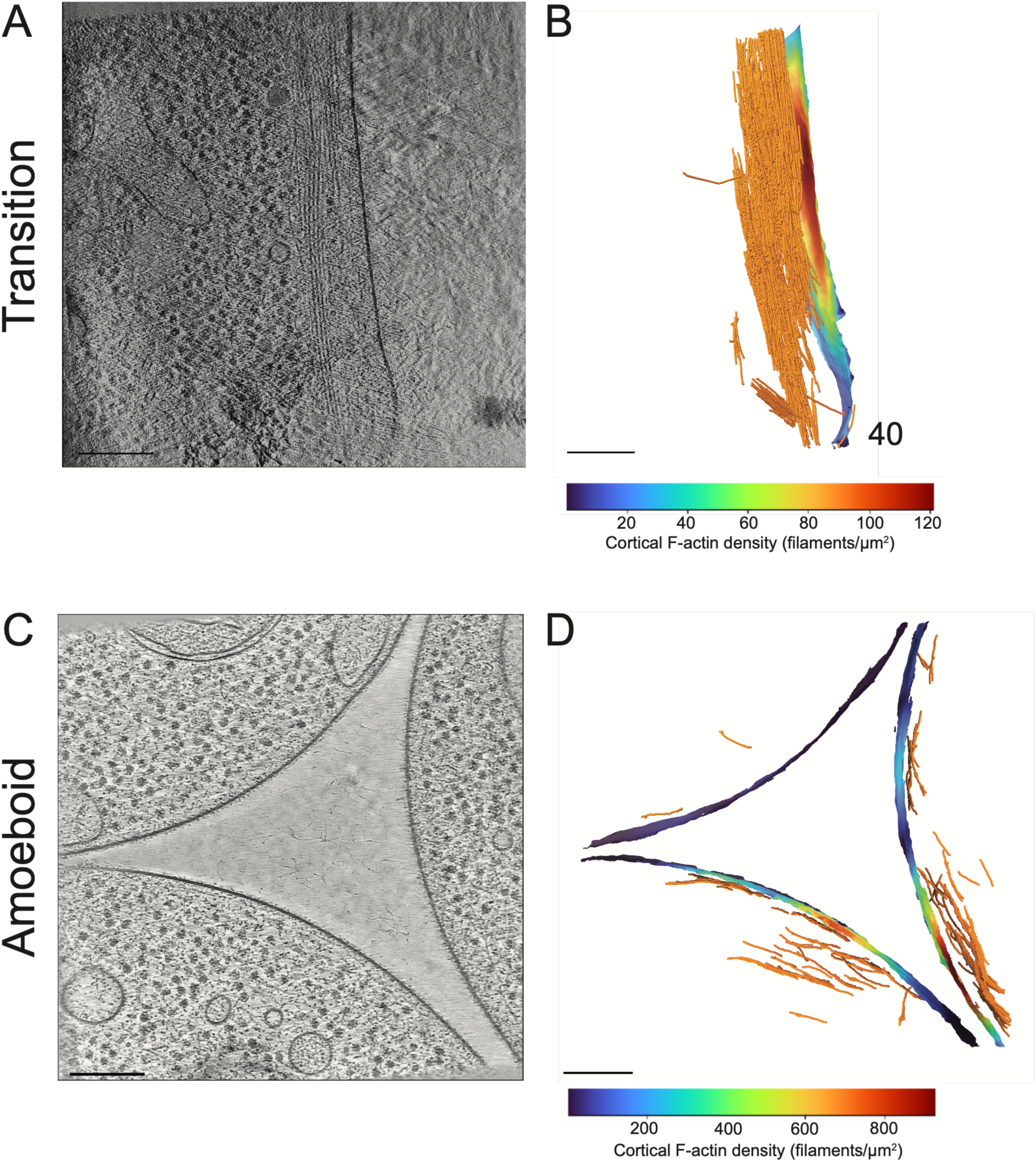
Quantification of cortical F-actin density. (A-B) Transition state. (C-D) Amoeboid state (note that the image shows three cells in close proximity). (A) and (C) are individual sections of reconstructed tomograms. (B) and (D) are segmented tomograms, with actin filaments depicted in orange, and the density of cortical filaments underneath the plasma membrane color-coded.

## List of supplementary movies

**Supplementary Movie 1. Live imaging reveals fast F-actin cortex formation under confinement.** Timelapse transmitted light/epifluorescence imaging of an *S. rosetta* strain expressing the live actin marker LifeAct-mCherry. Left panel: pink: F-actin, grey: transmitted light. Middle panel: transmitted light. Right panel: F-actin. Note onset of confinement at 17 s and near-instantaneous cortex formation at 19 s.

**Supplementary Movie 2. Live imaging reveals concomitant microvillar reabsorption and F-actin cortex formation.** Timelapse transmitted light/epifluorescence imaging of an *S. rosetta* strain expressing the live actin marker LifeAct-mCherry. Left panel: pink: F-actin, grey: transmitted light. Middle panel: transmitted light. Right panel: F-actin. Note onset of confinement at 21 s and gradual microvillar resorption, correlated with cortex formation, until 80 s.

**Supplementary Movie 3. Confinement induces a Ca^2+^ signaling pulse.** Timelapse transmitted light/epifluorescence imaging of an *S. rosetta* strain expressing the calcium signalign marker RGECO. Left panel: color scale: RGECO, grey: transmitted light. Middle panel: transmitted light. Right panel: RGECO. The color scale is the same as in Fig. 2.

**Supplementary Movie 4. Ionomycin treatment induces a Ca^2+^ signaling pulse.** Timelapse transmitted light/epifluorescence imaging of an *S. rosetta* strain expressing the calcium signalign marker RGECO. Left panel: transmitted light. Right panel: RGECO. Note the brief blebbing (from 78 to 111 sec) after ionomycin addition.

**Supplementary Movie 5. Ionomycin treatment induces blebbing.** Timelapse transmitted light/epifluorescence imaging of a wild-type *S. rosetta* strain. Ionomycin addition at 13 seconds.

**Supplementary Movie 6. Ionomycin treatment induces simultaneous microvillar reabsorption and F-actin cortex formation.** Timelapse transmitted light/epifluorescence imaging of an *S. rosetta* strain expressing the live actin marker LifeAct-mCherry. Left panel: transmitted light. Right panel: F-actin.

**Supplementary Movie 7. Ionomycin treatment induces myosin II condensation into cortical foci and fibers.** Timelapse transmitted light/epifluorescence imaging of an *S. rosetta* strain expressing the fluorescent myosin II marker MRLC-mTFP. Left panel: transmitted light. Right panel: myosin II.

**Supplementary Movie 8. Control cells bleb under confinement.** Controls corresponding to the ruthenium red-treated sample (Supp. Movie 9). Timelapse transmitted light/epifluorescence imaging of a wild-type *S. rosetta* strain.

**Supplementary Movie 9. Ruthenium red-treated cells show reduced blebbing under confinement.** Timelapse transmitted light/epifluorescence imaging of a wild-type *S. rosetta* strain.

**Supplementary Movie 10. Reconstructed and segmented cryo-tomogram of apical microvilli in the flagellate state.** Color code is the same as in Fig. 5.

**Supplementary Movie 11. Reconstructed and segmented cryo-tomogram of cortical microtubules in the flagellate state.** Color code is the same as in Fig. 5.

**Supplementary Movie 12. Reconstructed and segmented cryo-tomogram of cortical F-actin bundles in the transition state.** Color code is the same as in Fig. 5.

**Supplementary Movie 13. Reconstructed and segmented cryo-tomogram of retracting microvilli in the transition state.** Color code is the same as in Fig. 5.

**Supplementary Movie 14. Reconstructed and segmented cryo-tomogram of the persistent microtubule-organizing center in the amoeboid state.** Color code is the same as in Fig. 5.

**Supplementary Movie 15. Reconstructed and segmented cryo-tomogram of the F-actin cortex in the amoeboid state.** Color code is the same as in Fig. 5.

